# Inverted logic of ecDNA localization converts nuclear periphery into oncogenic transcription hubs in cancer

**DOI:** 10.64898/2026.09.11.751063

**Authors:** Yanbo Wang, Xiaowei Yan, Natasha E. Weiser, Ivy Tsz-Lo Wong, Shu Zhang, Yung-Hsin Huang, Rui Li, Katerina Kraft, Aditi Gnanasekar, Jun Tang, Cy Chittenden, Lotte Brückner, Imran Noorani, Charles Swanton, Anton G. Henssen, Andrew B. Stergachis, Nicolas Altemose, Howard Y. Chang, Paul S. Mischel

## Abstract

Extrachromosomal DNA (ecDNA) is a major driver of cancer pathogenesis, however, its spatial organization in the nucleus is not currently understood. Here we show that ecDNAs, in contrast to chromosomal DNA, preferentially localize to the nuclear periphery yet resist peripheral gene silencing. Single-molecule multi-omic sequencing and imaging-based CRISPR screening demonstrate enhanced ecDNA hub formation, H3K27ac and H3K9me3 bivalency, late DNA replication, long-range CpG hypomethylation, and increased transcriptional output when ecDNAs localize to the nuclear periphery. EcDNA anchors to the nuclear lamina through H3K9me3-dependent interactions with the Lamin B Receptor (LBR), which if interrupted, results in inward migration of ecDNA particles, reduced ecDNA congregation, and decreased transcription of ecDNA-encoded genes. These results reveal inherent advantages of the spatial organization of ecDNA that are not available to chromosomal DNA, enhancing ecDNA congregation and amplifying oncogenic transcriptional output, by localizing to the nuclear periphery of cancer cells.

## INTRODUCTION

Extrachromosomal DNA (ecDNA) is a prevalent mechanism of oncogene amplification in human cancers and a major driver of tumor evolution, intratumoral heterogeneity and therapeutic resistance^1–5^. These megabase-sized circular DNA elements harbor oncogenes together with regulatory sequences and reside outside the canonical chromosomal architecture^6,7^. A fundamental yet poorly understood property of ecDNA is its spatial organization within the nucleus. In most eukaryotic cells, nuclear positioning of DNA is closely linked to gene regulation: transcriptionally silent, heterochromatin-rich domains are preferentially localized to the nuclear periphery, whereas transcriptionally active domains are enriched in the nuclear interior^8,9^. EcDNA contains highly accessible chromatin and active histone marks, and is consequently highly transcribed^6,7,10^. However, ecDNA is frequently detected at the nuclear periphery^3,7,10–12^, raising the possibility that ecDNA may violate the canonical principles of genome organization and evade nuclear-peripheral repression through currently unknown mechanisms.

Here, we address the following fundamental questions: (1) How are ecDNA molecules spatially organized within the nucleus of a cancer cell? (2) What molecular mechanisms govern ecDNA spatial organization? And (3) how does ecDNA spatial organization regulate its oncogenic functions? We hypothesized that ecDNA possesses unique epigenomic and structural features that enable sustained transcriptional activity at the nuclear periphery, and that peripheral localization confers additional functional advantages to ecDNA. To test this hypothesis, we developed single-molecule multi-omic sequencing technologies that simultaneously profile nuclear position and multiple epigenetic modalities of individual ecDNA molecules, overcoming two major limitations of existing sequencing approaches that lack spatial information and cannot resolve ecDNA molecular heterogeneity^13^. Together with high-content microscopy, we elucidate the molecular principles that bridge ecDNA spatial organization to oncogenic functions (**Figure 1A**).

**Figure 1.**
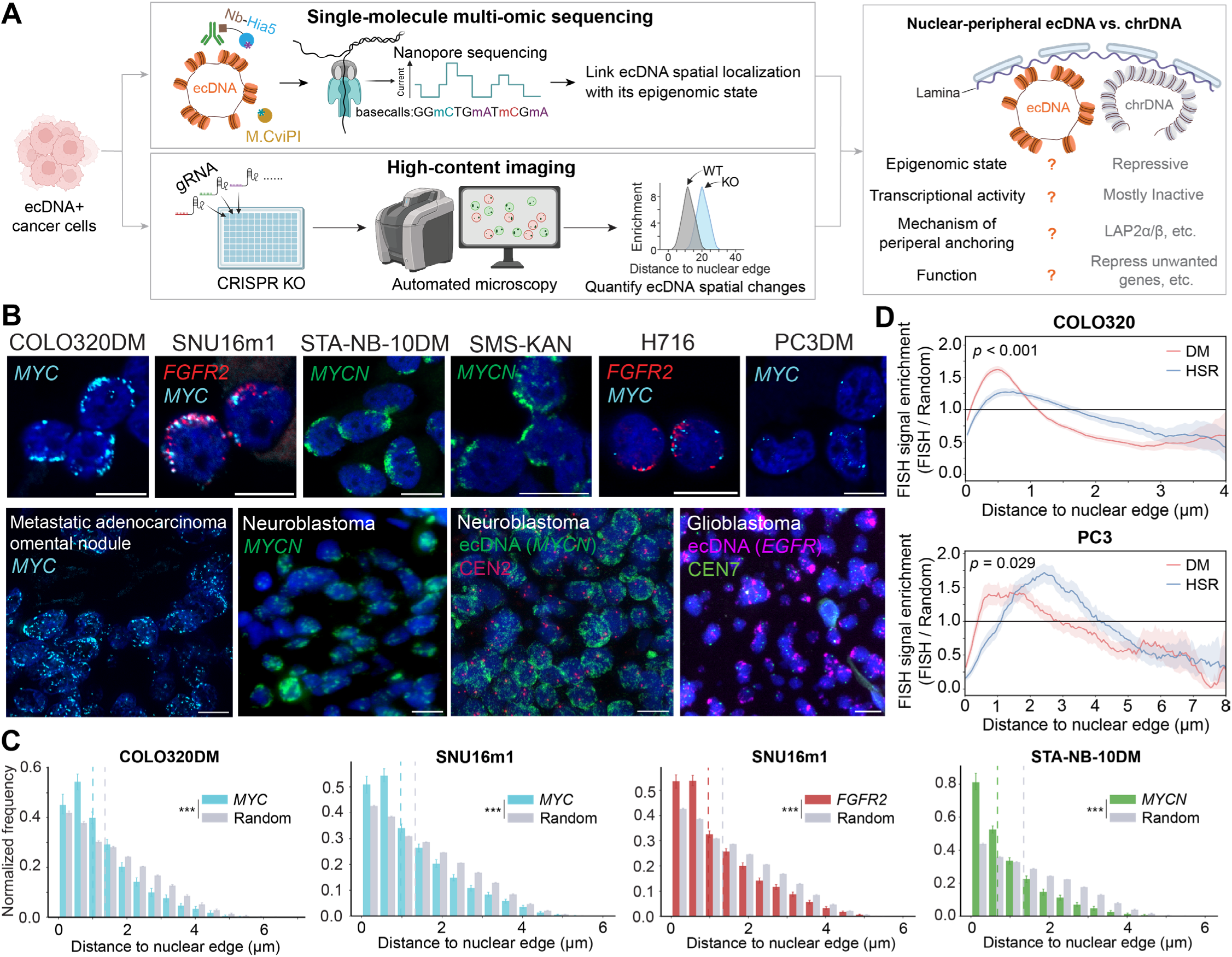
EcDNA preferentially localizes to the nuclear periphery in multiple cancer cell lines and patient samples of diverse cancer types. (A) Roadmap of this study. We established single-molecule multi-omic sequencing approaches to directly link ecDNA spatial localization with its chromatin state, revealing epigenomic features of nuclear-peripheral ecDNA that are distinct from chromosomal DNA. In parallel, we combined high-content imaging with targeted genetic perturbations to identify key factors that control ecDNA spatial localization. These complementary strategies converge to reveal the molecular mechanisms and the functional consequences of ecDNA spatial organization. (B) Representative DNA FISH images visualizing ecDNA molecules in cancer cells and patient tissue samples of diverse cancer types. COLO320DM and H716, colorectal cancer; SNU16m1, gastric cancer; STA-NB-10DM and SMS-KAN, neuroblastoma; and PC3DM, prostate cancer. Blue color represents DAPI-stained nuclei. For cancer cell lines, images acquired at the mid-nuclear plane are shown; for tissue samples, maximum-intensity projections are shown. Scale bar represents 10 μm. Scale bars in Neuroblastoma images are estimated. CEN2, centromere of chr2. CEN7, centromere of chr7. (C) Radial distribution of ecDNA distances to the nuclear edge, quantified from the DNA FISH data in Figure 1B, and compared with a random distribution in which ecDNA FISH signals were uniformly distributed across the nucleus. Dashed lines indicate the median of each distribution. Error bars represent 95% confidence intervals. *** represents *p* < 0.001. *P* values were calculated using a two-sided permutation test (5000 permutations) comparing the observed and random radial distributions. See Methods “Radial Distribution Analysis” for details. (D) FISH signal enrichment distributions of *MYC* DNA FISH signals of COLO320DM and PC3DM, and compared with the isogenic HSR lines (COLO320HSR and PC3HSR). To compute the FISH signal enrichment curves (FISH/Random), the radial distribution profile of FISH signals was divided by the corresponding random distribution. Shaded regions represent 95% confidence intervals. *P* values were computed using a two-sided permutation test (2000 permutations).

## RESULTS

### EcDNA remains active at nuclear periphery

We set out to visualize ecDNA and its location inside the nucleus by performing DNA fluorescence *in situ* hybridization (FISH) in cancer cell lines and patient tissue samples across different cancer types, including colorectal adenocarcinoma, prostatic adenocarcinoma, gastric carcinoma, neuroblastoma and glioblastoma (**Figure 1B**), revealing that ecDNA was preferentially localized to the nuclear periphery in these samples (**Figure 1C** and **Figure S1A**). When comparing oncogene DNA FISH signals between ecDNA-containing cell lines and their isogenic counterparts with the oncogene amplified on chromosomal homogeneously staining regions (HSR), we found that the oncogenes on ecDNA were more frequently localized to the nuclear periphery than their chromosomal HSR counterparts (**Figure 1D**). These observations demonstrate that nuclear-peripheral localization of ecDNA bearing different oncogenes can be detected in different types of cancer cells, including in bona fide clinical tumor samples.

To assess whether ecDNA remains active at nuclear periphery, we measured the DNA accessibility of nuclear-peripheral ecDNA given that DNA accessibility is highly correlated with transcriptional activity^14^. Existing bulk methods such as ATAC-seq cannot distinguish between peripheral versus interior ecDNA molecules. To fill this technical gap, we developed **lamin B1 DINO-seq**, a single-molecule sequencing technique that simultaneously measures DNA accessibility and nuclear spatial localization of individual DNA molecules, by integrating lamin B1 <u>Di</u>MeLo-seq^15,16^ with <u>NO</u>Me-seq^17–19^ (**Figure 2A**). In lamin B1 DINO-seq, permeabilized cells were treated with lamin B1 antibody and antibody-binding nanobody fused with the nonspecific deoxyadenosine methyltransferase Hia5 (Nb-Hia5)^16^. The lamin B1 antibody recruits Nb-Hia5 to the nuclear lamina, enabling Nb-Hia5 to selectively methylate adenines on lamina-associated DNA upon supplying S-adenosyl-methionine (SAM)^15^ (**Figure 2A**). Next, the GpC methyltransferase M.CviPI was applied to the cells, and only cytosines in the GC sequence context that are accessible to the enzyme were methylated^17^ (**Figure 2A**). Finally, genomic DNA was extracted and sequenced by base-modification-sensitive long-read sequencing (**Figure 2A**). Base modifications on each sequenced DNA molecule contain three types of molecular information (**Figure 2B**): adenine methylation (mA) provides spatial information about whether the DNA molecule is associated with the nuclear lamina^15^; cytosine methylation in the HCG sequence context (HmCG, where H represents A, C or T, and mC represents 5-methylcytosine) indicates the endogenous CpG methylation level; and cytosine methylation in the GCH context (GmCH) provides chromatin accessibility information of the DNA molecule^17^. Cytosines in GCG contexts were excluded from analysis because methylation at these sites cannot be unambiguously attributed to either DNA accessibility or endogenous CpG methylation^17^. Reads with mA levels above a defined threshold were classified as lamina-associated domain (LAD) reads, and those below were classified as non-LAD reads (**Figure 2B** and **Figure S1B**; see Methods). We found the LAD read enrichment is highly correlated with lamin B1 ChIP-seq signals, and the identified LAD reads enriched at the previously annotated constitutive lamina associated domains (cLAD) and depleted at the constitutive inter-lamina associated domains (ciLAD)^20^ (**Figure S1C** and **Figure S1D**), confirming the specificity of the LAD read classification.

**Figure 2.**
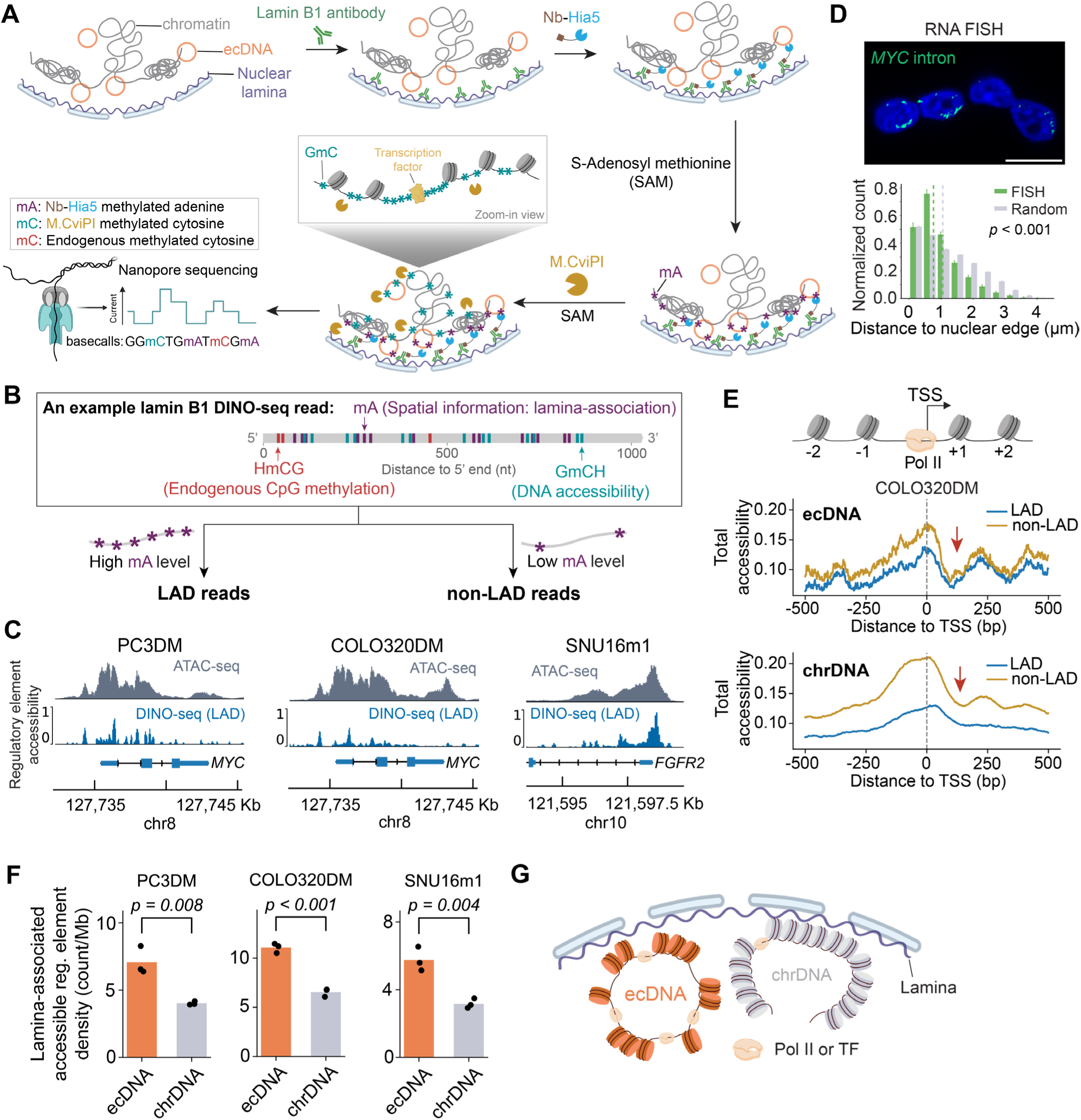
EcDNA remains active at nuclear periphery. (A) Schematic of lamin B1 DINO-seq. In lamin B1 DINO-seq, cells were permeabilized and treated with lamin B1 antibody, which binds to nuclear lamina. Next, antibody-binding nanobody fused with the nonspecific deoxyadenosine methyltransferase Hia5 (Nb-Hia5) was bound to lamin B1 antibody, and any unbound Nb-Hia5 was washed away. Cells were then incubated in a buffer containing SAM to activate adenine methylation in the vicinity of nuclear lamina. The GpC methyltransferase M.CviPI was applied into the cells, and only cytosines in the GC context that were accessible to the enzyme were methylated (i.e., accessible GC was methylated to GmC). Finally, genomic DNA was extracted and sequenced by base-modification-sensitive long-read sequencing, enabling simultaneous detection of methylated adenines (mA) deposited by Nb-Hia5, methylated cytosines (mC) in the GCH sequence context (GmCH) introduced by M.CviPI, and endogenous mC in the HCG sequence context (HmCG). H represents A, C or T. Cytosines in GCG contexts were excluded from analysis because methylation at these sites cannot be unambiguously attributed to either M.CviPI treatment or endogenous CpG methylation. (B) An example read from lamin B1 DINO-seq. Base modification mA (dark purple), GmCH (turquoise), HmCG (red) are labeled on the read. Lamin B1 DINO-seq reads were classified as a LAD read or non-LAD read based on its adenine methylation (mA) level. (C) ATAC-seq tracks and pseudo-bulk regulatory element accessibility tracks of LAD reads aligned to oncogene promoter regions in PC3DM, COLO320DM, and SNU16m1 cells. (D) Representative image of RNA FISH against *MYC* intron regions in COLO320DM cells, and radial distribution of the RNA FISH signals distances to the nuclear edge compared to all nuclear pixels (Random). Dashed lines represent median values of corresponding distributions. Error bars represent 95% confidence intervals. *P* values were computed using a two-sided permutation test (5000 permutations), comparing the observed difference in distributions. Scale bar represents 10 μm. (E) DNA accessibility analysis of lamin B1 DINO-seq reads at TSS in COLO320DM cells. Line profiles show the total accessibility (i.e., the sum of regulatory element accessibility and linker DNA accessibility) of lamin B1 DINO-seq reads at each position around TSS. Red arrows indicate the +1 nucleosome footprints. (F) Comparison of accessible regulatory element density on LAD ecDNA and LAD chrDNA reads. The density was calculated as the total number of accessible regulatory elements detected on LAD ecDNA or LAD chrDNA reads divided by the corresponding cumulative read length of those reads. Dot on the bar plot represents individual replicates (n=3). *P* values were calculated using two-sided Student’s t-test. (G) Schematic depicting a model in which nuclear-peripheral ecDNA has a higher density of accessible regulatory elements than nuclear-peripheral chrDNA. These accessible regulatory elements may be occupied by transcription factors (TF) and RNA polymerase II (Pol II).

To analyze DNA accessibility, we generated the pseudo-bulk GmCH/GCH track by computing, at each genomic position, the fraction of reads carrying a GmCH modification at the position (**Figure S2A**). Comparison of the pseudo-bulk GmCH/GCH track with ATAC-seq revealed GmCH enrichment at ATAC-seq peaks, but also substantial GmCH signal outside the ATAC-seq peak regions, resulting in a relatively high background in the pseudo-bulk GmCH/GCH track (**Figure S2B**). This is because both accessible regulatory elements (e.g., enhancers and promoters) and linker DNA connecting adjacent nucleosomes are accessible to M.CviPI, and are therefore methylated by the enzyme^18^ (**Figure S2C**). Consequently, the observed GmCH signals reflect the sum of **regulatory element accessibility** and **linker DNA accessibility**. Only accessible regulatory elements indicate nucleosome-depleted open chromatin^21^. To obtain regulatory element accessibility for each read, we applied Density-Based Spatial Clustering of Applications with Noise (DBSCAN)^22^ to cluster GmCH modifications on each read and selectively remove short-range GmCH clusters corresponding to linker DNA (**Figure S2B**; see Methods). This approach leverages the fact that linker DNA is typically <85 bp in length, substantially shorter than accessible regulatory elements^21,23^, thereby enabling measurement of regulatory element accessibility from the GmCH modifications on each read (**Figure S2B**). After removing linker DNA signals, the pseudo-bulk regulatory element accessibility track (i.e., the fraction of reads carrying an accessible regulatory element at each genomic position) was highly correlated with ATAC-seq (**Figure S2B** and **Figure S2D**), and exhibited sharper peak profiles owing to the near single-base-pair resolution of DINO-seq (**Figure S2B**). These data demonstrate that the lamin B1 DINO-seq faithfully captures DNA accessibility information of individual DNA molecules.

To test whether the oncogenes remain active on nuclear-peripheral ecDNA molecules, we performed lamin B1 DINO-seq on previously established monoclonal ecDNA-positive cell lines, including PC3DM (prostatic adenocarcinoma)^10^, COLO320DM (colorectal adenocarcinoma)^6^, and SNU16m1 (gastric carcinoma)^7^. In these models, the vast majority of amplicon DNA copies reside on ecDNA^6,7,10^, indicating that reads aligned to the ecDNA amplicon predominantly originate from ecDNA molecules (hereafter referred to as ecDNA reads). Lamin B1 DINO-seq revealed regulatory element accessibility peaks at oncogene promoters and enhancers on LAD ecDNA reads, indicating that ecDNA remains accessible even at the nuclear periphery (**Figure 2C**). Consistently, nascent mRNA from the ecDNA-harboring oncogene was preferentially detected near the nuclear periphery, confirming oncogene RNA transcription activities on nuclear-peripheral ecDNA molecules (**Figure 2D**). By centering lamin B1 DINO-seq reads at transcription start sites (TSS) (**Figure S3A**), we assessed DNA accessibility of LAD and non-LAD reads in the vicinity of TSS (**Figure 2E**). We observed well-positioned +1 nucleosome footprints for both LAD and non-LAD ecDNA molecules across different cell lines (**Figure 2E** and **Figure S3B**). The well-positioned +1 nucleosome indicates the assembly of the preinitiation complex at TSS^24^. In contrast, the footprint of well-positioned +1 nucleosome was not observed for LAD chromosomal DNA (chrDNA) molecules (**Figure 2E** and **Figure S3B**). These data suggest that unlike LAD chrDNA, LAD ecDNA are primed for transcription. Finally, by calculating the average density of accessible regulatory elements on individual LAD reads, we found that LAD ecDNA harbored a significantly higher density of accessible regulatory elements than LAD chrDNA (**Figure 2F**). Together, these data demonstrate that unlike chrDNA, ecDNA molecules at nuclear periphery remain highly active (**Figure 2G**).

### EcDNA is enriched for both repressive and active epigenomic features

To investigate how ecDNA remains transcriptionally active at the nuclear periphery, we examined its associated histone modifications. H3K9 trimethylation (H3K9me3) is a hallmark of constitutive heterochromatin and is classically associated with transcriptional repression at the nuclear periphery^20^. Concurrent *MYC* DNA FISH and H3K9me3 immunofluorescence (IF) in COLO320DM cells confirmed strong enrichment of H3K9me3 signals at the nuclear periphery (**Figure 3A**). Unexpectedly, nuclear-peripheral ecDNA exhibited H3K9me3 IF intensities comparable to its surrounding peripheral chrDNA (**Figure 3A**), suggesting that ecDNA itself may also be highly decorated with H3K9me3. Consistent with this observation, both CUT&RUN and ChIP-seq showed pronounced H3K9me3 enrichment across the ecDNA amplicon (**Figure 3B** and **Figure S4A**). When comparing the H3K9me3 CUT&RUN signals to copy-number-normalized (CN-normalized) ATAC-seq signals, we observed that H3K9me3 and accessible chromatin domains generally occupied distinct megabase-scale regions (**Figure 3B**). However, ecDNA represented a striking exception: both H3K9me3 and CN-normalized ATAC-seq signals were enriched at the ecDNA amplicon (**Figure 3B**). Notably, more than one quarter of ATAC-seq peaks on ecDNA lie within 1 kb of an H3K9me3 peak, over fourfold higher than on chrDNA (**Figure 3C**), indicating ecDNA amplicon is enriched for repressive and active chromatin features in close genomic proximity.

**Figure 3.**
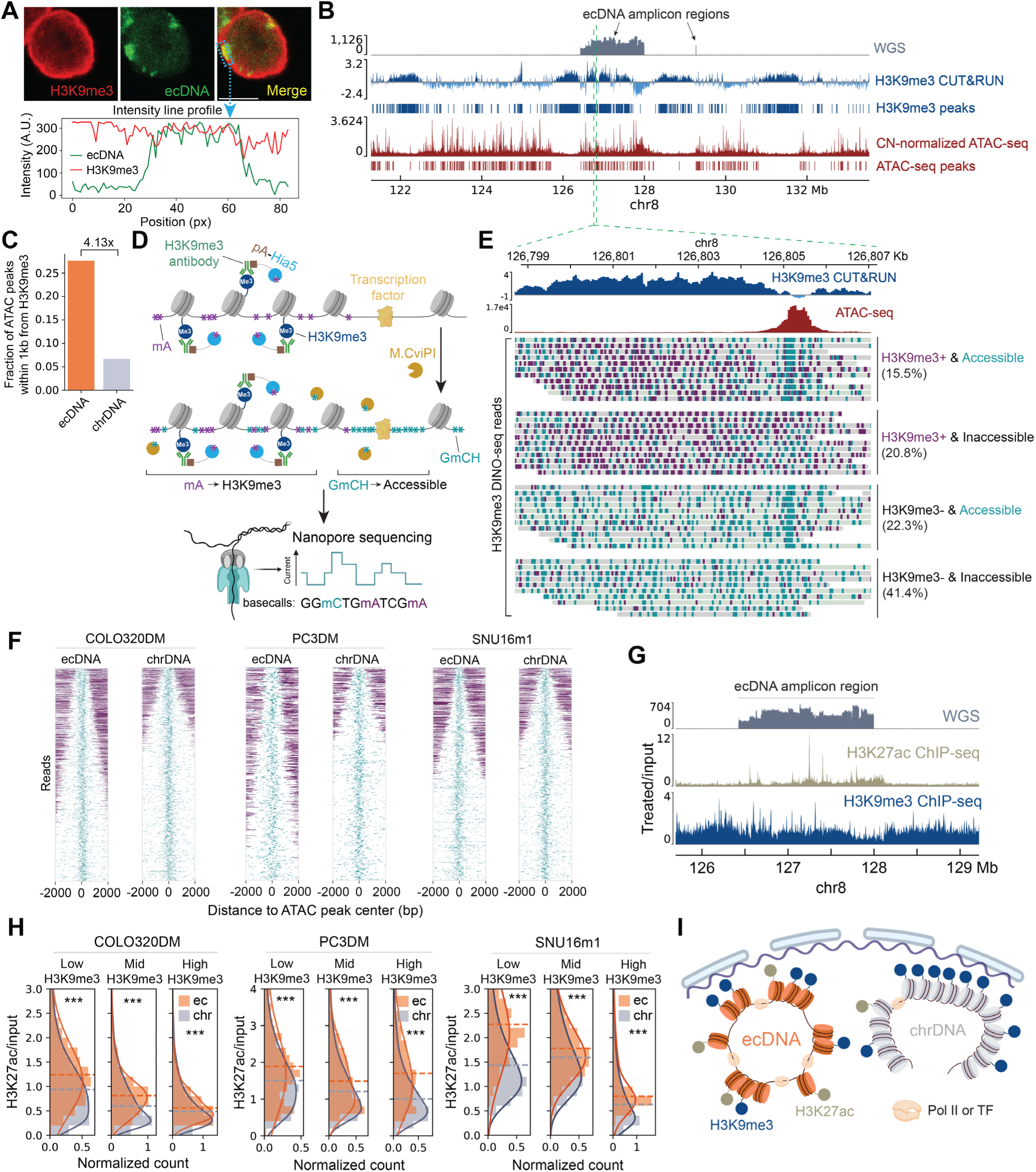
EcDNA is enriched for both repressive and active epigenomic features (A) Representative image of concurrent *MYC* DNA FISH (to visualize ecDNA) and H3K9me3 IF in COLO320DM cells. The light-blue dashed rectangle highlights a region where the DNA FISH and H3K9me3 IF signals overlap; intensity line profiles of the DNA FISH and IF signals across this region are shown. These data suggest that H3K9me3 levels on nuclear-peripheral ecDNA are comparable to those of the surrounding chromosomal DNA. Scale bar represents 10 μm. (B) Whole genome sequencing (WGS), H3K9me3 CUT&RUN, copy-number (CN)-normalized ATAC-seq tracks of COLO320DM cells. H3K9me3 and ATAC-seq peaks are indicated using vertical bars below each track. (C) Comparison of fractions of ATAC-seq peaks that are within 1kb from H3K9me3 peaks between ecDNA amplicon regions and chrDNA. (D) Schematic of H3K9me3 DINO-seq. The H3K9me3 antibody recruits pA-Hia5 to H3K9me3 nucleosomes, enabling pA-Hia5 to selectively methylate adenines surrounding the H3K9me3 nucleosomes. M.CviPI was applied to the cells, and only cytosines in the GC sequence context that are accessible to the enzyme were methylated. Finally, genomic DNA is sequenced by modification-sensitive, long-read sequencing. (E) Example H3K9me3 DINO-seq reads aligned to a genomic locus within the ecDNA amplicon, where H3K9me3 CUT&RUN signals and an ATAC-seq peak are immediately next to each other. Base modification mA (dark purple) and GmCH (turquoise) were labeled on individual reads. Each read is displayed with a grey or light-green background, indicating alignment to the plus or minus strand of the reference genome, respectively. The H3K9me3 DINO-seq reads were classified into four groups based on their H3K9me3 level and whether they had an accessible regulatory element overlapping the ATAC-seq peak. Percentages of reads classified into each group are shown. Green dashed lines show the zoomed-out location of this genomic locus in the ecDNA amplicon in Figure 3B. (F) H3K9me3 DINO-seq reads are centered at ATAC-seq peaks and sorted based on the distance between an accessible regulatory element (turquoise) and its nearest H3K9me3 signal (dark purple) on each read. The top 5%, 5% and 2% ranked reads for COLO320DM, PC3DM and SNU16m1 are shown. ChrDNA reads were randomly subsampled to match the ecDNA read counts. (G) Tracks of WGS, H3K27ac ChIP-seq and H3K9me3 ChIP-seq from COLO320DM cells. (H) Comparison of H3K27ac levels of ecDNA and chrDNA at high, middle and low H3K9me3 genomic regions where ecDNA and chrDNA had comparable H3K9me3 levels. See Methods for definitions of High, Mid and Low H3K9me3 regions. Dashed lines indicate the median values of the corresponding histograms. Solid curves represent kernel density estimates (KDE) of the signal distributions. *** represents *p* < 0.0001. *P* values were computed using two-sided Mann–Whitney U tests. (I) EcDNA exhibits simultaneous enrichment of both repressive and active marks.

Bulk ATAC-seq and CUT&RUN (**Figure 3B**) cannot distinguish whether the co-enrichment of open chromatin and H3K9me3 on ecDNA arises from molecular heterogeneity (i.e., distinct “active” versus “inactive” ecDNA molecules) or from bivalency on individual ecDNA molecules (i.e., one ecDNA molecule simultaneously carries repressive and active chromatin marks in vicinity to each other). To resolve this question, we performed **H3K9me3 DINO-seq**, which simultaneously profiles H3K9me3 histone modification and DNA accessibility on the same DNA molecule (**Figure 3D**). The workflow of H3K9me3 DINO-seq was identical to lamin B1 DINO-seq, except that an H3K9me3 antibody was used in place of the lamin B1 antibody, and protein A-fused Hia5 (pA-Hia5) was used instead of Nb-Hia5 to achieve higher affinity for the H3K9me3 antibody^15^ (**Figure 3D**). In H3K9me3 DINO-seq, mA indicates H3K9me3 histone modification, and GmCH provides DNA accessibility information (**Figure 3D**). H3K9me3 signal and accessible regulatory elements on each read were identified by analyzing mA and GmCH modifications on the read using DBSCAN (**Figure S4B**). The pseudo-bulk H3K9me3 signal track from the DINO-seq showed nearly identical enrichment patterns to CUT&RUN and ChIP-seq data (**Figure S4A**), confirming the specificity of H3K9me3 detection of DINO-seq.

H3K9me3 DINO-seq revealed distinct epigenomic landscapes on different ecDNA molecules (**Figure 3E**). For instance, at an ecDNA amplicon locus where bulk H3K9me3 and ATAC-seq peaks lie in proximity (**Figure 3E**), individual ecDNA molecules were classified into four distinct epigenomic states depending on whether they contained H3K9me3 signals and/or were accessible at the ATAC-seq peak position. Remarkably, 15.5% of ecDNA molecules were highly accessible at the ATAC-seq peak position immediately adjacent to H3K9me3 signals (**Figure 3E**), demonstrating that a single ecDNA molecule can simultaneously carry repressive and active epigenomic features in close proximity. When reads were sorted based on the distance between an accessible regulatory element and its nearest H3K9me3 signal, ecDNA displayed a substantially higher fraction of reads with accessible regulatory elements surrounded by H3K9me3 compared to chrDNA across different cell lines (**Figure 3F**). These results suggest that individual ecDNA molecules are more likely to harbor repressive and active epigenomic features in close proximity than chrDNA.

Given that H3K9me3-marked chrDNA is typically compacted into heterochromatin^25,26^, we reasoned that ecDNA might counteract H3K9me3-associated compaction by concurrently enriching histone modifications associated with chromatin decompaction. We therefore hypothesized that ecDNA molecules not only accumulate H3K9me3 but also retain high levels of H3K27ac, a histone mark associated with chromatin decompaction^27,28^. Supporting this idea, ChIP-seq showed both H3K9me3 and H3K27ac enrichment across the ecDNA amplicon, in contrast to the adjacent chrDNA region where only H3K9me3 was present (**Figure 3G**). Quantification of H3K9me3 and H3K27ac ChIP-seq signals in 1-kb bins across the genome further showed that ecDNA carried overall higher levels of both histone modifications compared with chrDNA (**Figure S4C**). Importantly, at genomic regions where ecDNA and chrDNA had comparable H3K9me3 levels, ecDNA consistently exhibited higher H3K27ac (**Figure 3H**). Together, these data indicate that ecDNA is co-enriched for repressive and active histone modifications (**Figure 3I**).

### Lamin B receptor tethers ecDNA at nuclear periphery

To identify mechanisms driving ecDNA’s peripheral localization, we developed an optical screening platform that integrates CRISPR-mediated gene knockout (KO) with quantitative DNA FISH in COLO320DM cells, to identify genes that regulate ecDNA localization in the nucleus (**Figure 4A**). Although DNA FISH enables direct visualization of ecDNA, it is typically low throughput and subject to technical variability, limiting robust quantitative spatial analysis. To overcome these limitations, we optimized DNA FISH for a 96-well imaging plate format, enabling cost-effective, scalable sample preparation and image acquisition while preserving sufficient cell numbers to capture ecDNA’s intrinsic cell-to-cell heterogeneity. To control for technical variability across wells, we incorporated an internal spike-in strategy in which mCherry-labeled cells nucleofected with Cas9-sgRNA (i.e., KO cells) were mixed 1:1 with mock nucleofected (no sgRNA) GFP-labeled control cells and co-seeded into the same wells (**Figure 4A**). Because fluorescent protein signals are lost during FISH, cell identity (i.e., KO or control) was preserved through sequential imaging: fluorescent protein markers were imaged before FISH, ecDNA signals were imaged afterward, and the two imaging datasets were computationally aligned to match cell identity with ecDNA FISH signals at single-cell resolution (**Figure 4A**).

**Figure 4.**
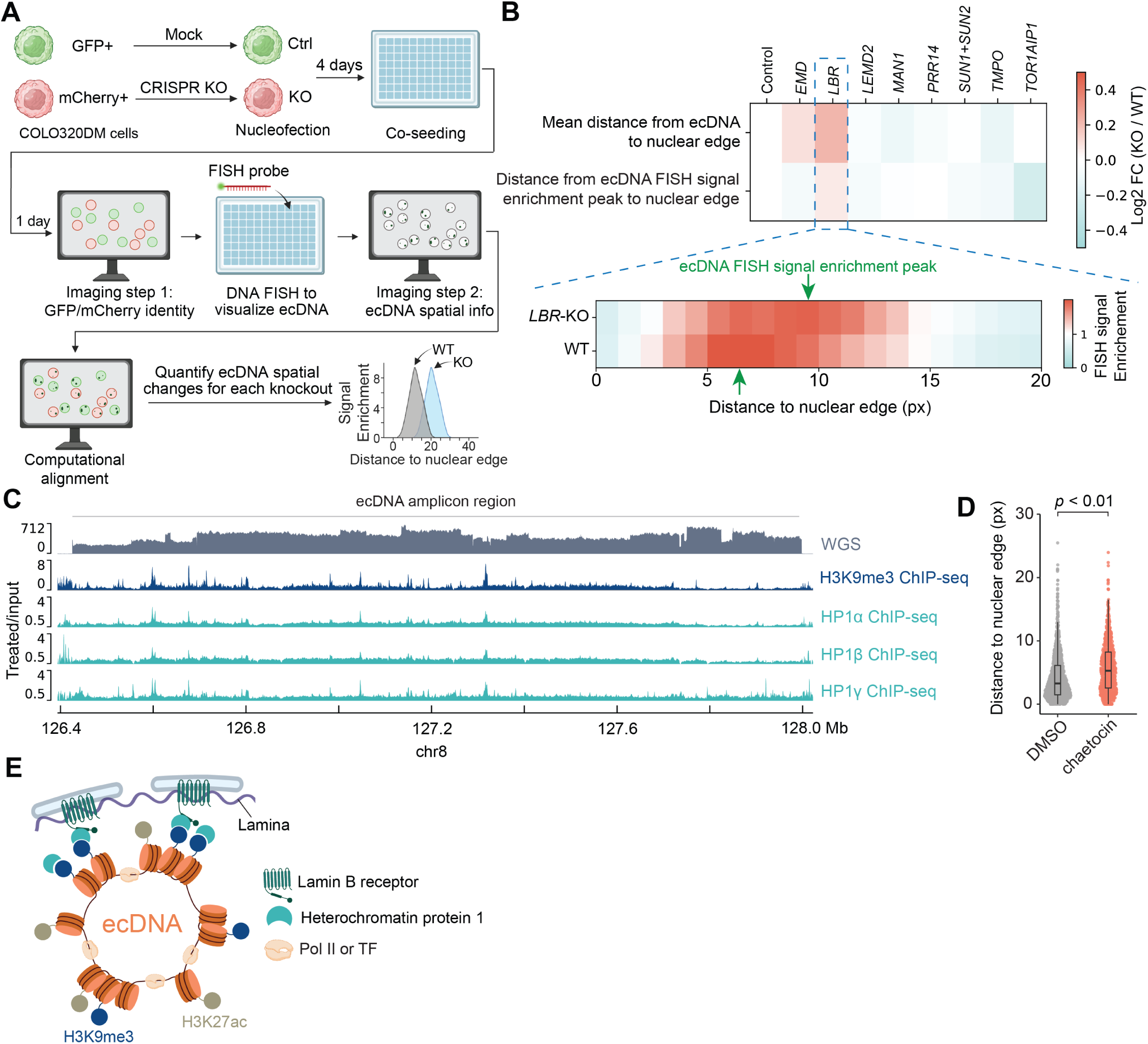
EcDNA is tethered at nuclear periphery through H3K9me3-dependent, *LBR*-mediated interactions. (A) Schematic of the optical screening platform. CRISPR knockout (KO) nucleofection was performed in COLO320DM cells expressing mCherry, while mock nucleofection (no sgRNA) was performed in COLO320DM cells expressing GFP as a control (Ctrl). Four days later, KO (mCherry⁺) and Ctrl (GFP⁺) cells were mixed and co-seeded into imaging plates. After 24 h, cells were fixed on the imaging plates, the identity of each cell (mCherry⁺ KO or GFP⁺ Ctrl) was recorded. DNA FISH was then performed to visualize ecDNA, and the resulting images were computationally aligned to match cell identity with ecDNA FISH signals at the single-cell level. Finally, ecDNA spatial organization was quantified and compared between mCherry⁺ KO and GFP⁺ Ctrl cells. (B) Quantification of ecDNA spatial changes following knockout of each gene in the screen. The top heatmap summarizes ecDNA spatial changes after each knockout using two metrics: mean distance from ecDNA to nuclear edge, and distance from ecDNA FISH signal enrichment peak to the nuclear edge. Mean ecDNA distance to the nuclear edge was calculated by computing the mean FISH-intensity-weighted distance of ecDNA to the nuclear edge in each cell, then averaging these values across cells. EcDNA FISH signal enrichment was computed by dividing the radial distribution profile of ecDNA FISH signals by the corresponding random distribution, as in Figure 1D. The ecDNA FISH signal enrichment peak indicates the position at which the ecDNA FISH signal is most enriched after normalizing to the random distribution. The bottom shows ecDNA FISH signal enrichment distribution heatmaps for *LBR*-KO and WT cells. Green arrows indicate the ecDNA FISH signal enrichment peak positions for *LBR*-KO and WT cells. Pixel size (px) is 0.11 µm. (C) WGS, H3K9me3 ChIP-seq and HP1 ChIP-seq tracks at the ecDNA amplicon region in COLO320DM cells. (D) Quantification of ecDNA spatial changes in COLO320DM cells treated with 100 nM chaetocin for 1 day. Each dot represents FISH-intensity-weighted mean ecDNA distance to the nuclear edge in a single cell. Pixel size (px) is 0.11 µm. Boxes represent the interquartile range (IQR) with center lines indicating the median; whiskers extend to 1.5× IQR. *P* value was calculated using a two-sided Kolmogorov-Smirnov test. (E) EcDNA is decorated with H3K9me3 histone modifications, which recruits HP1, a known binding partner of the inner nuclear membrane protein lamin B receptor.

Utilizing this optical screening platform, we tested a list of genes encoding proteins primarily located at nuclear periphery with the potential to associate with DNA^29–31^ (**Figure 4B**). For each gene, a pool of three sgRNAs was used to ensure efficient knockout. The screen revealed that loss of lamin B receptor (LBR) (**Figure S5A**) caused the most striking relocalization of ecDNA toward the nuclear interior (**Figure 4B**). Consistently, ChIP-seq data revealed that heterochromatin protein 1 (HP1), a known interaction partner of the lamin B receptor^32^, was recruited to H3K9me3-decorated ecDNA^33^ (**Figure 4C**). Treating the cells with chaetocin (an inhibitor of histone methyltransferases for H3K9me3)^34^ reduced H3K9me3 levels on ecDNA (**Figure S5B**) and led to significant displacement of ecDNA away from the nuclear periphery (**Figure 4D**). Together, these results suggest that H3K9me3-dependent, *LBR*-mediated interactions tether ecDNA to the nuclear lamina (**Figure 4E**).

### EcDNA nuclear-peripheral localization is associated with long-range CpG hypomethylation and late replication

We next investigated the functional consequences associated with ecDNA positioning at the nuclear periphery. Using lamin B1 DINO-seq (**Figure 2A**), we compared endogenous CpG methylation landscapes between nuclear-peripheral and nuclear-interior ecDNA molecules (**Figure 5A**). We observed long-range cytosine hypomethylation in the HCG sequence context for LAD ecDNA reads as compared to that for non-LAD ecDNA reads (**Figure 5A**). Such long-range hypomethylation has previously been linked to incomplete epigenetic maintenance associated with late DNA replication^35^. This observation led us to hypothesize that nuclear-peripheral ecDNA has delayed DNA replication compared to interior ecDNA.

**Figure 5.**
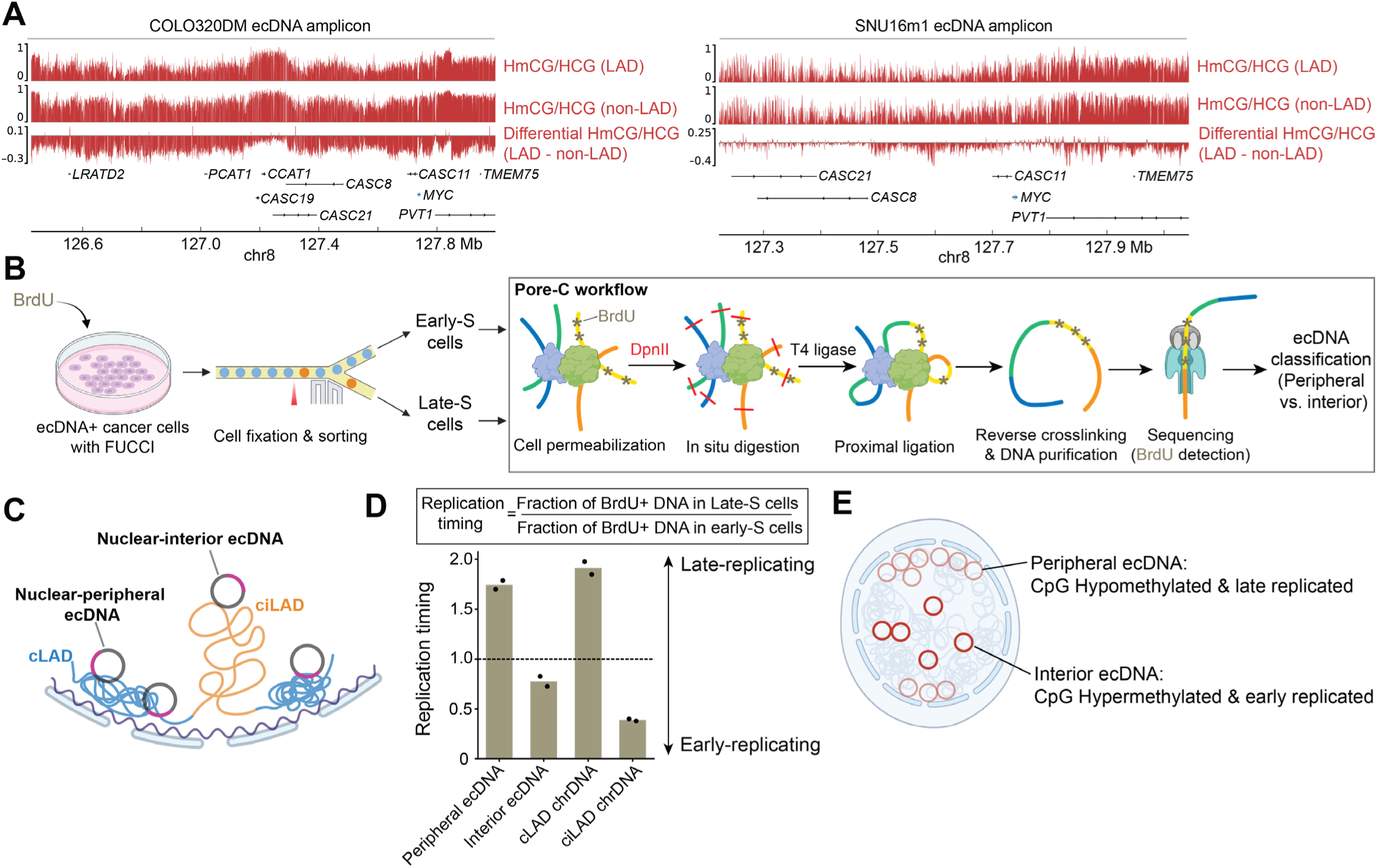
Nuclear-peripheral ecDNA exhibits long-range CpG hypomethylation and delayed DNA replication. (A) Comparison of cytosine methylation in the HCG context between LAD and non-LAD ecDNA reads in COLO320DM and SNU16m1 cells. For each sample, the first and second tracks are the pseudo-bulk HmCG/HCG tracks (i.e., fraction of cytosines methylated in the HCG context at each genomic position) for LAD and non-LAD ecDNA reads of lamin B1 DINO-seq, respectively. The third track shows the difference between the LAD and non-LAD methylation profiles (Differential HmCG/HCG; LAD subtracts non-LAD), where the predominant negative values indicate long-range hypomethylation for LAD reads as compared to non-LAD reads. H denotes A, C or T. (B) Schematic of the single-molecule assay to compare replication timing of nuclear-peripheral and - interior ecDNA. (C) Each Pore-C read aligned to the ecDNA amplicon was classified as originating from nuclear-peripheral or nuclear-interior ecDNA based on whether it contacted predominantly cLAD or ciLAD chrDNA, respectively. (D) Quantification of replication timing of nuclear-peripheral ecDNA, nuclear-interior ecDNA, cLAD chrDNA and ciLAD chrDNA. Dots indicate individual replicates. (E) EcDNA nuclear-peripheral localization is associated with long-range CpG hypomethylation and late replication.

To test this hypothesis, we established a single-molecule assay that jointly measures replication timing and nuclear localization for individual ecDNA molecules (**Figure 5B**). We engineered the fluorescent ubiquitination-based cell cycle indicator (FUCCI)^36,37^ into COLO320DM cells and pulse-labeled replicating DNA with BrdU, followed by cell sorting to isolate early-S and late-S populations based on FUCCI fluorescence (**Figure 5B**). Sorted cells proceeded for Pore-C^38^ to detect both BrdU incorporation^39^ and chromatin-contact information for each DNA molecule, in which permeabilized nuclei were digested with DpnII, proximally ligated, and sequenced using Nanopore (**Figure 5B** and **Figure S6A**). For each Pore-C read aligned to the ecDNA amplicons, we determined whether it originated from nuclear-peripheral or nuclear-interior ecDNA based on whether the read contacted predominantly cLAD or ciLAD chrDNA (**Figure 5C** and **Figure S6B**; see Methods). Because BrdU marks newly synthesized DNA, we evaluated the replication timing of nuclear-peripheral and-interior ecDNA reads by calculating the fraction of BrdU-positive ecDNA reads in early-S and late-S cells (**Figure 5D**). Nuclear-peripheral ecDNA showed a higher fraction of BrdU-positive reads in late-S than in early-S cells, indicating late replication (**Figure 5D**). In contrast, nuclear-interior ecDNA exhibited the opposite pattern, with a higher fraction of BrdU-positive reads in early-S cells, indicating earlier replication timing (**Figure 5D**). As internal controls, cLAD and ciLAD chrDNA behaved as expected for late-and early-replicating domains, respectively^40^ (**Figure 5D**). The replication timing difference between interior and peripheral ecDNA may explain the previous observation of disorganized ecDNA replication^41^. These results support our hypothesis that nuclear-peripheral ecDNA is associated with long-range hypomethylation and delayed DNA replication compared with nuclear-interior ecDNA (**Figure 5E**).

### EcDNA nuclear-peripheral localization promotes large ecDNA hub formation and oncogene expression

EcDNA molecules tend to cluster into hubs, thereby enhancing intermolecular interactions and transcription^7,42,43^. The nuclear-peripheral architecture may facilitate hub formation by providing anchoring sites (**Figure 4E**) and by imposing geometric constraints that favor molecular clustering. We therefore hypothesized that ecDNA would form larger hubs near the nuclear periphery than those located in the nuclear interior. To test this hypothesis, we visualized ecDNA by DNA FISH and quantified the total FISH intensity of individual ecDNA FISH objects as a function of distance to the nuclear edge (**Figure 6A** and **Figure S7A**). Each ecDNA FISH object, defined as a continuous patch of ecDNA FISH signal, represents either an ecDNA hub or a single ecDNA molecule (**Figure 6A**). EcDNA FISH objects closer to the nuclear edge exhibited substantially higher FISH intensity than those farther from the nuclear edge (**Figure 6B**), consistent with our hypothesis that larger ecDNA hubs form at the nuclear periphery. Similar results were also observed in SNU16m1 cells, which harbor two distinct ecDNA species carrying different oncogenes^7^ (**Figure 6C**), as well as in a glioblastoma patient sample (**Figure S7B**). Concordantly, Pore-C analysis revealed that nuclear-peripheral ecDNA displayed significantly higher ecDNA-ecDNA contact frequencies compared with nuclear-interior ecDNA (**Figure S7C**). These data demonstrate markedly elevated ecDNA congregation at the nuclear periphery compared with the nuclear interior.

**Figure 6.**
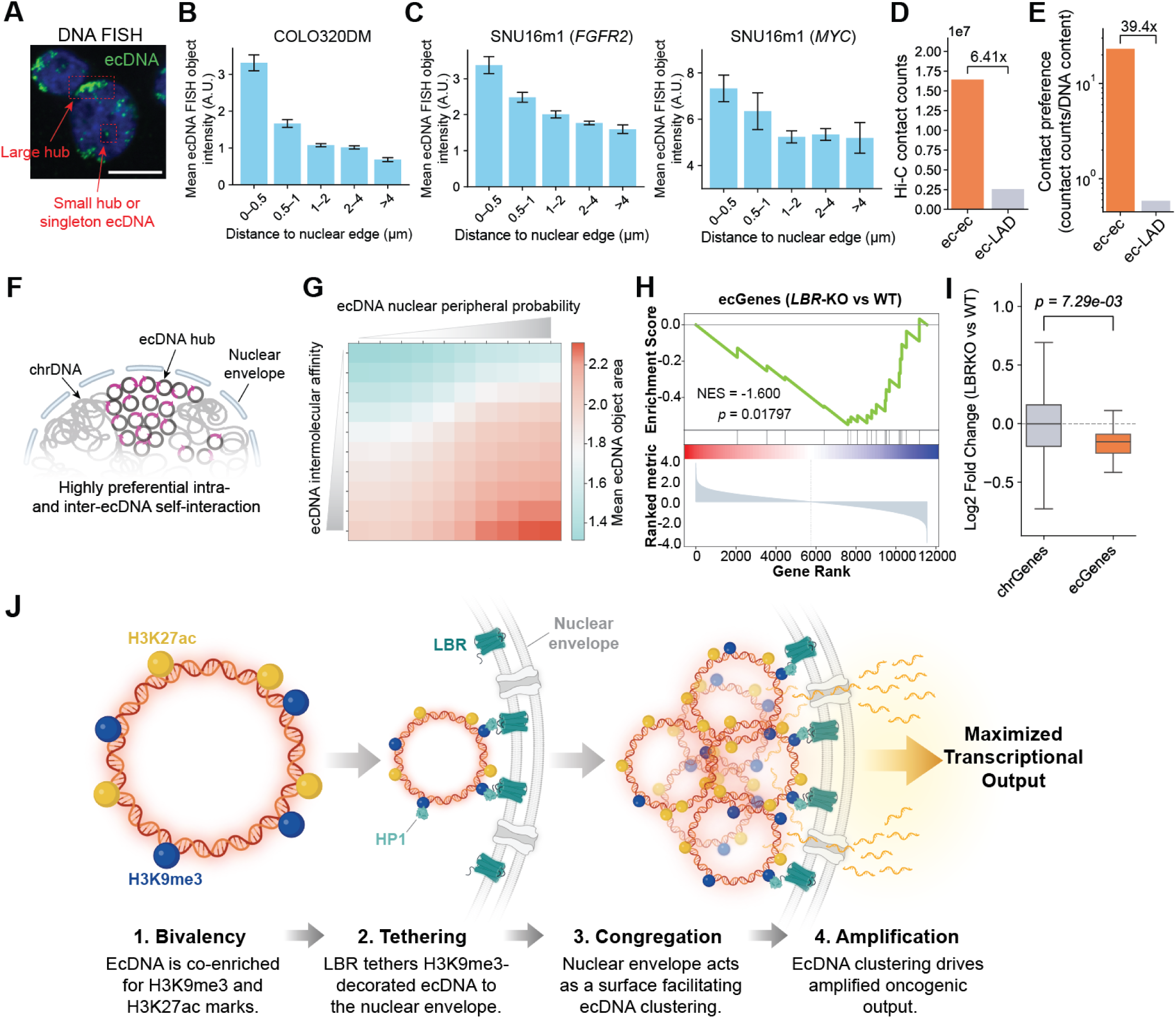
Nuclear lamina anchoring promotes large ecDNA hub formation and enhances transcriptional output of ecDNA-encoded genes (A) Representative *MYC* DNA FISH image of COLO320DM cells. Red dashed rectangles highlight ecDNA FISH objects corresponding to a large ecDNA hub at nuclear periphery, and a small ecDNA hub/singleton ecDNA at nuclear interior. Blue color represents DAPI-stained nuclei. Scale bar represents 10 μm. (B) Bar plots of mean FISH intensity of individual ecDNA FISH objects as a function of distance to the nuclear edge in COLO320DM cells. Error bars indicate standard error of the mean. (C) Bar plots of mean FISH intensity of individual ecDNA FISH objects as a function of distance to the nuclear edge for the two ecDNA species carrying *FGFR2* and *MYC* oncogenes, respectively, in SNU16m1 cells. Error bars indicate standard error of the mean. (D) Comparison of total Hi-C contact counts between ecDNA-ecDNA interactions and ecDNA-LAD chrDNA interactions in COLO320DM cells. LAD regions on chrDNA were inferred from Lamin B1 ChIP-seq, with LADs called in genomic bins where log2(treated/input) > 0 (Figure S1C). (E) Comparison of Hi-C contact preferences between ecDNA-ecDNA interactions and ecDNA-LAD chrDNA interactions. Contact preference was calculated by normalizing the total Hi-C contact counts between ecDNA-ecDNA or ecDNA-LAD chrDNA (Figure 6D) by their respective cellular DNA content (Figure S7D). (F) EcDNA at the nuclear periphery shows a strong bias toward intra-and inter-ecDNA interactions rather than interactions with LAD chrDNA, thereby establishing a locally active chromatin environment at the nuclear periphery. (G) Heatmap showing the mean ecDNA object area as a function of ecDNA intermolecular affinity and ecDNA nuclear peripheral probability in simulated nuclei. (H) Gene set enrichment analysis of ecGenes following *LBR* knockout in COLO320DM cells. Normalized enrichment score (NES) and nominal *p*-value are shown. (I) Differential expression levels of ecDNA-encoded genes (ecGenes) and chromosomal DNA-encoded genes (chrGenes) after *LBR* KO in COLO320DM cells. Each dot represents an ecGene. Boxes represent the IQR with center lines indicating the median; whiskers extend to 1.5× IQR. *P* values were calculated using two-sided Mann–Whitney U tests. (J) EcDNA hijacks nuclear-peripheral architecture to enhance oncogenic function. EcDNA is co-enriched for H3K9me3 and H3K27ac and maintains a decompacted, transcriptionally permissive state. The nuclear envelope imposes a two-dimensional spatial constraint that promotes ecDNA congregation into large transcriptional hubs at the nuclear periphery, thereby amplifying the transcriptional output of ecDNA-encoded genes. LBR, lamin B receptor. HP1, heterochromatin protein 1.

Given that most ecDNA molecules localize to the nuclear periphery (**Figure 1B**), we assessed contact preferences between ecDNA and LAD chrDNA. Hi-C analysis revealed a strong bias toward intra-and inter-ecDNA self-interaction: contact counts between ecDNA regions were 6.4-fold higher than contacts between ecDNA and LAD chrDNA regions (**Figure 6D**), even though whole-genome sequencing revealed that LAD chrDNA contributes sixfold more cellular DNA content than ecDNA (**Figure S7D**), indicating a nearly 40-fold enrichment of intrahub ecDNA-ecDNA contact preference relative to ecDNA-LAD chrDNA contacts (**Figure 6E**). These data suggest that ecDNA preferentially engages in intra-and inter-ecDNA interactions rather than interactions with LAD chrDNA, thereby establishing a locally active chromatin environment at the nuclear periphery^9^ (**Figure 6F**). Finally, to assess whether ecDNA nuclear-peripheral localization facilitates larger hub formation, we performed *in silico* simulations to generate nuclei containing the same number of ecDNA copies while systematically varying ecDNA peripheral localization probability and ecDNA-ecDNA intermolecular affinity (**Figure S7E**). We found that increasing either of these factors individually, or both in combination, led to an increase in the average size of ecDNA hubs (**Figure 6G**). Collectively, these results demonstrate that ecDNA nuclear-peripheral localization promotes the formation of large ecDNA hubs.

Given that ecDNA hubs promote intermolecular interactions and transcription^7^, we hypothesized that disrupting interactions between ecDNA and the nuclear membrane would reduce ecDNA congregation and consequently decrease the expression of genes carried on ecDNA. Consistent with this hypothesis, DNA FISH analysis revealed that *LBR*-knockout (*LBR*-KO) in COLO320DM cells exhibited a reduced frequency of large ecDNA hubs compared with wild-type (WT) cells (**Figure S7F**). Notably, disruption of ecDNA nuclear-peripheral localization by *LBR* knockout led to a preferential reduction in the expression of ecDNA-encoded genes (ecGenes, **Figure 6H** and **Figure 6I**), without altering average ecDNA copy number (**Figure S7G**). These results suggest that nuclear-peripheral localization of ecDNA enhances ecGene expression.

Finally, we examined the expression level of *LBR* in patient samples from The Cancer Genome Atlas (TCGA). We stratified TCGA samples by ecDNA status using AmpliconArchitect^44^ and compared *LBR* expression level between ecDNA-positive and ecDNA-negative tumors (**Figure S7H**). We found that *LBR* was significantly upregulated in ecDNA-positive tumors (**Figure S7H**). In addition, genes encoding several H3K9me3 histone methyltransferases and HP1 were also significantly upregulated in ecDNA-positive tumors compared to ecDNA-negative tumors (**Figure S7I**). These data suggest potential functional importance of LBR and the H3K9me3 methyltransferase machinery in ecDNA-positive cancers.

## DISCUSSION

### EcDNA hijacks nuclear-peripheral architecture to enhance oncogenic function

One of the most distinctive features of ecDNA is that it is not constrained by canonical chromosome territories; Indeed, ecDNA’s unique epigenomic landscape enables it to exploit nuclear-peripheral architecture to amplify its transcriptional output (**Figure 6J**). Although heterochromatin is canonically localized to the nuclear periphery, ecDNAs have inverted this logic. Mechanistically, ecDNA molecules enriched for H3K9me3 engage the inner nuclear membrane protein lamin B receptor, driving their preferential accumulation at the nuclear periphery (**Figure 6J**). Lamina acting as ecDNA docking sites, together with the two-dimensional geometric constraints imposed by the nuclear envelope, potentially transforming the random collision of ecDNAs from three-dimensional to two-dimensional search, increases the probability of ecDNA-ecDNA contacts and promotes the formation of ecDNA hubs. Despite being tethered to a heterochromatic environment, ecDNA remains decompacted and transcriptionally active, with enrichment of accessible regulatory elements and H3K27ac. The enhanced congregation of ecDNA at the periphery facilitates intermolecular enhancer-gene interactions, thereby further increasing oncogene expression^7^. Together, our data illustrate how cancer cells harness the unique spatial and epigenomic properties of ecDNA to amplify ecGene expression.

We demonstrate the ability of ecDNA to maintain transcriptional activity despite lamina association and H3K9me3 exposure. This behavior cannot be attributed simply to the fact that ecDNA selectively amplifies promoters that are intrinsically resistant to LAD repression^9,45,46^. Evidence from comparing ecDNA-containing lines with their isogenic HSR counterparts (each isogenic pair have highly similar amplified sequences) shows that ecDNA exhibits substantially higher transcriptional activity^10^ while occupying the nuclear periphery more frequently than HSR (**Figure 1D**), suggesting that mechanisms beyond the primary sequence underlie ecDNA’s resistance to LAD repression. Our data support a model in which ecDNA molecules establish a locally transcriptionally active environment at the nuclear periphery, where individual ecDNA molecules are mostly surrounded by other ecDNAs, fostering primarily ecDNA-ecDNA interactions, rather than ecDNA-LAD chrDNA interactions (**Figure 6D**), creating an active chromatin environment that insulates ecDNA from LAD repression^9^ (**Figure 6F**).

### Spatial and epigenetic dependencies of ecDNA expose potential vulnerabilities

We identify LBR as the key molecular anchor that actively tethers ecDNA molecules to the nuclear periphery. While LBR is known to mediate peripheral chromatin anchoring in some cell types during development^47,48^, its role in cancer gene expression remains largely unclear. We show that ecDNA hijacks LBR’s chromatin binding activity to scaffold the oncogenic transcription hub at nuclear periphery, enhancing expression of genes encoded on ecDNA. Notably, *LBR* was recently reported to be upregulated in metastatic melanoma and associated with decreased patient survival, where cancer cells exploit LBR’s sterol reductase activity to enhance nuclear deformability and promote metastatic invasion^49^. We also revealed that *LBR* is upregulated in ecDNA-containing tumors (**Figure S7H**). These findings emphasize the functional significance of LBR in cancer and nominate it as a candidate therapeutic target.

The recognition that nuclear-peripheral positioning of ecDNA is associated with late DNA replication and increased transcription also provides important mechanistic insights into ecDNA evolution. The elevated transcription of ecDNA intensifies transcription-replication conflicts, leading to replication stress and slow replication forks^10^. Their late replication timing could further increase mutation rates by reducing the available window for resolving stalled replication forks and repairing DNA before cell division^50,51^. Consistent with this model, ecDNA is enriched for the SBS8 mutational signature^52^, which predominantly arises from late-replication errors^53^. Thus, nuclear-peripheral localization may accelerate ecDNA evolution and contribute to the increased susceptibility of ecDNA-positive cancers to therapies targeting DNA replication stress^10^.

Furthermore, because CpG methylation represents a relatively stable epigenetic component that can be faithfully propagated through cell divisions^54^, the distinct CpG methylation landscapes observed between nuclear-peripheral and nuclear-interior ecDNA (**Figure 5A**) suggest that ecDNA nuclear positioning may be heritable across generations. For example, nuclear-peripheral ecDNA molecules in parental cells may remain at the nuclear periphery in daughter cells. Such heritable spatial organization could stabilize intermolecular interactions among different ecDNA species and facilitate the coordinated inheritance of fitness-enhancing ecDNA species combinations^3^.

Finally, lamina-associated DNA double-strand breaks (DSBs) are predominantly repaired through canonical or alternative non-homologous end joining (c-NHEJ or alt-NHEJ)^55^. The preferential association of ecDNA with lamina, together with its elevated DNA damage^10^, suggests that ecDNA-positive cancers may be particularly dependent on c-NHEJ or alt-NHEJ pathways for survival. Consistent with this notion, depletion of alt-NHEJ components has been shown to reduce ecDNA copy number in cancer cells^43^. Moreover, because ecDNA is co-enriched for H3K27ac and H3K9me3 and forms large transcriptional hubs at nuclear periphery, combinatorial targeting of these epigenetic programs (e.g., JQ1^7^ and chaetocin) may achieve synergistic effects by concurrently disrupting ecDNA spatial organization and oncogene transcription. Together, these findings highlight yet another way that the altered spatial organization of ecDNA in the nucleus could contribute to tumor pathogenesis. Future systematic screening of small molecules targeting these pathways will be instrumental in uncovering selective vulnerabilities and developing effective therapeutic strategies for ecDNA-driven cancers.

### DINO-seq resolves ecDNA heterogeneity in spatial and epigenomic context

EcDNA exhibits a high degree of molecular heterogeneity, posing a fundamental technical challenge to its systematic characterization. Existing sequencing approaches lack spatial resolution and cannot link multiple epigenetic modalities on the same DNA molecule^13^, making it difficult to dissect how ecDNA spatial organization interfaces with its epigenetic state. To overcome this technical gap, we developed a suite of single-molecule multi-omic sequencing technologies. Specifically, we developed DINO-seq to simultaneously profile protein binding, DNA accessibility, and CpG methylation on individual ecDNA molecules (**Figure 2A** and **Figure 3D**). In addition, we repurposed Pore-C as a spatially-resolved modification-sensitive sequencing platform, and interrogated ecDNA replication timing and proximity contacts as a function of nuclear position (**Figure 5B**). These approaches provide a generalizable framework for directly linking multiple epigenetic layers on single DNA molecules in their spatial context. Furthermore, the long-read nature of these technologies makes them particularly well suited to resolving complex genomic architectures and elucidating the epigenomic principles governing gene fusions^56^, repetitive elements^57^, and chromatin bivalency^58,59^ at unprecedented resolution in cancer and developmental biology. We anticipate that our single-molecule multi-omic strategy will be broadly applicable for studying genome organization and function, epigenetic landscapes, and regulatory mechanisms^60^.

## Resource availability Lead contact

Further information and requests for resources and reagents should be directed to and will be fulfilled by the lead contact, Howard Y. Chang and Paul S. Mischel.

## Materials availability

This study did not generate new unique reagents.

## Data and code availability

Sequencing data generated in this study will be deposited in the NCBI Gene Expression Omnibus (GEO) under the accession number GSE319725. Microscopy data generated in this study will be deposited on Figshare. Custom code used in this study will be made available on Github.

## Acknowledgement

We thank the members of the Chang and Mischel labs for discussion. We would like to thank all the patients for their generosity in participating in the clinical trials and giving permission to share their data. Glioblastoma tissue samples were obtained from the University Hospital Southampton NHS Foundation Trust as part of BRAIN UK, which is supported by Brain Tumor Research and has been established with the support of the British Neuropathological Society and the Medical Research Council. This project was supported by the Cancer Grand Challenges partnership funded by Cancer Research UK (CGCATF-2021/100012 and CGCATF-2021/100025) and the National Cancer Institute (OT2CA278688 and OT2CA278635) to P.S.M. and H.Y.C.. Y.W. is a Schmidt Science Fellow. X.Y. is a Damon Runyon Fellow supported by the Damon Runyon Cancer Research Foundation (DRG-2474-22). N.E.W. was supported by the National Cancer Institute (K08CA296931). N.A. is an HHMI Hanna H. Gray Faculty Fellow, a Pew Biomedical Scholar, and a Biohub Investigator. H.Y.C. was an Investigator of the Howard Hughes Medical Institute.

## Author Contributions

Y.W., X.Y., H.Y.C. and P.S.M. conceived and designed the study. Y.W. performed DINO-seq, Pore-C, Nanopore WGS experiments and data analysis. Y.W. performed RNA-seq experiments. Y.W. and S.Z. analyzed RNA-seq assay data. S.Z. analyzed gene expression of TCGA data. X.Y., and N.E.W. performed DNA FISH experiments. X.Y. performed DNA FISH-IF, RNA FISH and optical screen experiments. X.Y. conducted simulations to generate nuclei with ecDNA. X.Y. and Y.W. performed imaging analysis. A.G. and J.T. supported microscopy and imaging analysis. I.T.-L.W. generated COLO320DM-FUCCI cell line, performed BrdU treatment and cell sorting for replication timing study. Y.W., N.E.W., and R.L. performed ChIP-seq experiments. N.E.W. performed CUT&RUN experiments and Y.W. performed CUT&RUN analysis. Y.W. and S.Z. conducted ChIP-seq data analysis. Y.-H.H. performed ATAC-seq experiments. Y.W. and S.Z. conducted ATAC-seq data analysis. Y.W., S.Z., and K.K. conducted Hi-C data analysis. N.A., C.C., and A.B.S. prepared reagents for DINO-seq and advised on nanopore sequencing experiments. L.B., I.N., A.G.H and C.S. provided DNA FISH images from patient samples and advised on the data analysis. H.Y.C. and P.S.M. supervised the studies. Y.W., X.Y., H.Y.C., and P.S.M. wrote the manuscript with input from all authors.

## Methods Cell culture

Human prostate cancer cell lines PC3DM and PC3HSR, and colorectal cancer cell lines COLO320DM and COLO320HSR, were cultured in Dulbecco’s Modified Eagle’s Medium (DMEM; Corning) supplemented with 10% fetal bovine serum (FBS; Gibco) and 1% penicillin–streptomycin. COLO320DM and COLO320HSR were purchased from ATCC, and PC3DM and PC3HSR were established as previously described^10^. SNU16m1 cells were derived from the parental SNU16 cell line as previously described^7^ and maintained in DMEM/F12 (Gibco, 11320-033) supplemented with 10% FBS and 1% penicillin–streptomycin. STA-NB10DM cells were cultured in RPMI-1640 medium (Gibco) supplemented with 1 % of penicillin–streptomycin, 10 % of fetal calf serum (FCS) (Thermo Fisher), 1 % Glutamax (Thermo Fisher), 1 % Sodium Pyruvat (Thermo Fisher) and 10 mM HEPES (Sigma). SMS-KAN cells were cultured in RPMI1640 medium (Gibco), supplemented with 10% FBS and 1% penicillin– streptomycin. To engineer monoclonal COLO320DM-FUCCI cells, parental COLO320DM cells were plated into a 6-well plate and transfected by preparing a mixture containing 1.6µg tFucci(CA)5 plasmid (Addgene, #153521), 0.4µg Super PiggyBac transposase expression vector (System Biosciences, PB210PA-1), 200µL Opti-MEM (Gibco), and 2µL X-tremeGENE^TM^ HP DNA Transfection Reagent (Roche, 6366236001). The component was mixed thoroughly by pipetting and incubated at room temperature for 15 mins. The mixture was added to the cells overnight and the medium was replaced the next day. Single COLO320DM-FUCCI cells were then plated into a 96-well plate for monoclonal expansion to generate monoclonal COLO320DM-FUCCI cells. All cell lines were maintained at 37 °C in a humidified incubator with 5% CO₂ and were routinely tested negative for mycoplasma contamination. For chaetocin treatment experiments, COLO320DM cells were incubated with 100nM chaetocin (MedChemExpress, HY-N2019) in the culture medium for 24 h at 37 °C prior to the imaging or ChIP-seq experiments.

### DINO-seq

Protein A-Hia5 was expressed and purified as previously described^15^, with a detailed protocol available online: dx.doi.org/10.17504/protocols.io.bv82n9ye. Anti-rabbit nanobody-Hia5 was purified as previously described^16^. Purification details of nanobody-Hia5 are available online: dx.doi.org/10.17504/protocols.io.6qpvrqbyzlmk/v1.

DINO-seq integrates the DiMeLo-seq protocol^15^ with NOMe-seq protocol^17^. One million cells were pelleted at 500g for 3 minutes and washed with PBS. Pelleted cells were resuspended in 1 ml of DW buffer (0.02% digitonin, 20 mM HEPES-potassium hydroxide buffer, pH 7.5, 150 mM sodium chloride, 0.5 mM spermidine, 1 Roche cOmplete EDTA-free tablet (Roche, 11873580001) per 50 ml buffer and 0.1% bovine serum albumin (BSA)), followed by a 5-minutes incubation on ice. The sample was spun down at 500 g for 3 minutes at 4 °C. The pellet was resuspended in TW buffer (0.1% Tween-20, 20 mM HEPES-potassium hydroxide, pH 7.5, 150 mM sodium chloride, 0.5 mM spermidine, 1 Roche cOmplete EDTA-free tablet per 50 ml buffer and 0.1% BSA), and spun down at 500 g for 3 minutes at 4 °C. The pellet was resuspended in TW buffer with 1:50 diluted primary antibody (LMNB1, Abcam, ab16048; H3K9me3, Active Motif, 39065), and incubated for 2 h at 4 °C on a rotator. The sample was pelleted at 500 g for 3 minutes at 4 °C, and washed twice with TW buffer with a 4 °C rotating incubation for 5 min between spins. The sample was then resuspended in TW buffer with 200 nM Hia5 construct (Nb-Hia5 for Lamin B1 DINO-seq, and pA-Hia5 for H3K9me3 DINO-seq), and incubated for 2 h at 4 °C on a rotator. The sample was pelleted at 500 g for 3 minutes at 4 °C, and washed twice with TW buffer with a 4 °C rotating incubation for 5 min between spins. The sample was resuspended in 100 µl of AB buffer (15 mM Tris, pH 8.0, 15 mM sodium chloride, 60 mM potassium chloride, 1 mM EDTA, pH 8.0, 0.5 mM EGTA, pH 8.0, 0.5 mM spermidine, 0.1% BSA and 800 µM S-adenosylmethionine (SAM)) and incubated at 37 °C for 2 h. The sample was gently resuspended every 30 minutes, and replenished SAM halfway through the 2-hour incubation by adding an additional 800 µM SAM. The sample was pelleted at 500 g for 3 minutes at 4 °C, and resuspended in 225 µl of NOMe buffer (1x GC Reaction Buffer (NEB, M0227S), 0.3 M sucrose, 96 μM SAM, 100 U M.CviPI (NEB, M0227S)). The sample was incubated for 30 minutes at 37 °C, and replenished SAM halfway through the 30-minute incubation by adding an additional 96 µM SAM. The sample was pelleted at 500 g for 3 minutes, and DNA was purified using the MagAttract HMW DNA Kit (Qiagen, 67563) according to the manufacturer’s protocol. Nanopore sequencing libraries were prepared using the Ligation Sequencing Kit V14 (Oxford Nanopore Technologies, SQK-LSK114) following the manufacturer’s instructions and sequenced on an Oxford Nanopore PromethION 24 platform.

### DINO-seq data analysis

#### Basecalling and alignment

Base calling of canonical bases and modified bases (6mA and 5mC) was performed using Dorado (v0.9.1) and models dna_r10.4.1_e8.2_400bps_sup@v5.0.0, dna_r10.4.1_e8.2_400bps_sup@v5.0.0_6mA@v1 and dna_r10.4.1_e8.2_400bps_sup@v5.0.0_5mC_5hmC@v1. Base called reads were aligned to hg38 reference genome using minimap2^63^ (v2.26-r1175).

#### Read filtering

By comparing the GmCH density of individual reads (i.e., the number of GmCH modifications on the read divided by aligned read length) from DINO-seq with a negative control experiment (i.e., DINO-seq without adding M.CviPI), we found a small fraction of reads have as minimal levels of GmCH density as the reads in the negative control experiment (**Figure S8A**). These could be due to incomplete permeabilization, or small number of cells clumped together, making cells buried inside inaccessible to M.CviPI. We therefore filtered out these reads with minimal GmCH density by setting a threshold as shown in **Figure S8A**. In addition to the reads with the minimal GmCH density, we also observed a small fraction of reads with very high GmCH density (**Figure S8B**). Because the footprint size of a nucleosome is ∼ 146 bp, and the size of a linker DNA between two nucleosomes is typically in the range of 0-80 bp^23^, the fraction of accessible DNA in a read that does not overlap with ATAC-seq peaks (i.e., the fraction of linker DNA in a read) should be less than 80/(146+80) = 0.35. The small fraction of extremely accessible reads was also observed in the previously reported single-molecule footprinting assays, and they are likely from dead cells^62^. Consistently, these reads with very high levels of GmCH do not show the nucleosome footprint signature when plotting the histogram of inaccessible DNA lengths on these reads, in contrast to the reads with the fraction of accessible DNA less than 0.35 (**Figure S8B**).

#### LAD read classification

For each read, the positions of methylated adenines (mA) on the read were detected from Nanopore sequencing, and the Density-Based Spatial Clustering of Applications with Noise (DBSCAN)^22^ in Scikit-learn (v1.5.1) was performed to identify mA signal clusters using parameters eps=75 and min_samples=2 given that the footprint of a nucleosome is generally larger than 75 bp^64^. Each mA signal cluster represents a lamina-associated linker DNA between adjacent nucleosomes. We then calculated the mA level of the read using total length of mA clusters not overlapping with any ATAC-seq peaks on the read divided by the alignment length of the read. We sorted all reads aligned to chromosomal DNA (i.e., regions outside ecDNA amplicons) based on their mA levels, and determined the mA level threshold where 5% of the reads are above the threshold. Reads with mA levels higher than the threshold are classified as LAD reads, and reads with mA levels lower than the threshold are classified as non-LAD reads. To generate the LAD read coverage in **Figure S1C**, LAD reads were extracted from the aligned bam of lamin B1 DINO-seq using pysam (v0.22.1) and computed coverage bigwig file using deepTools (v3.5.4.post1).

#### Regulatory element accessibility and total accessibility

To obtain the regulatory element accessibility for each read, the positions of methylated cytosines in GCH sequence context (GmCH) on the read were detected from Nanopore sequencing, and DBSCAN was performed to identify GmCH signal clusters using parameters eps=75 and min_samples=1 (**Figure S2B**). Each GmCH signal cluster represents either a linker DNA between adjacent nucleosomes or an accessible regulatory element. To remove GmCH signal clusters that come from linker DNA, we filtered out the signal clusters with length less than 85 bp (**Figure S2B**). The remaining GmCH signal clusters represent accessible regulatory elements. The pseudo-bulk regulatory element accessibility track was computed by calculating, at each genomic position, the fraction of reads carrying an accessible regulatory element overlapping with the position (**Figure S2B**). The density of lamina-associated accessible regulatory elements in **Figure 2F** was calculated as the total number of accessible regulatory elements detected in LAD-associated reads divided by the total length of LAD-associated reads mapped to either ecDNA amplicon regions or chromosomal DNA regions (i.e., the hg38 genome excluding ecDNA amplicon intervals). EcDNA amplicon coordinates for each cell line were obtained from AmpliconArchitect (v.1.3.r8) output as previously described^10^, with minor manual adjustments to refine amplicon boundaries based on WGS coverage profiles. Total accessibility (**Figure 2E** and **Figure S3B**) reflects the sum of regulatory element accessibility and linker DNA accessibility. Total accessibility at each genomic position was calculated as the fraction of reads harboring DBSCAN-clustered GmCH signals (i.e., include both accessible regulatory elements and linker DNA) overlapping that position.

#### H3K9me3 DINO-seq read processing

For each read, the positions of GmCH and mA on the read were detected from Nanopore sequencing. As shown in **Figure S4B**, the GmCH and mA modifications on each read were analyzed separately. GmCH signals were processed as described in the previous section to identify accessible regulatory elements. mA signals overlapping with any ATAC-seq peaks on the read were filtered out to avoid potential confounding from Hia5 methylation of nucleosome-depleted DNA neighboring H3K9me3-bound nucleosomes. Next, two rounds of DBSCAN clustering were performed because H3K9me3 domains typically span broad genomic regions. In the first round of DBSCAN using parameters eps=75 and min_samples=4, individual mA events were grouped into discrete mA clusters. In the second round of DBSCAN using parameters eps=1000 and min_samples=3, adjacent mA clusters were merged into extended H3K9me3 signals.

### ATAC-seq

Bulk ATAC-seq samples were prepared as previously described^65^, and the eluted DNA samples (∼20 µl) were used for subsequent library amplification following the OMNI-ATAC-seq protocol^66^. The libraries were sequenced on an Illumina NovaSeq 6000 platform with paired-end 150 bp reads. Raw reads were first QC and trimmed adapter by Trim Galore (v0.6.10). Adapter-trimmed reads were then aligned to the hg38 genome using bwa (v0.7.18). Duplicate fragments of aligned reads were removed using Picard (v2.27.5). Because the ecDNA amplicon region has substantially higher sequencing depth, which would reduce peak-calling *q*-values compared to chromosomal regions, we subsampled reads from the ecDNA amplicon according to ecDNA copy number (determined from WGS) to equalize sequencing depth between the ecDNA amplicon and chromosomal regions. The subsampled bam file was used for ATAC-seq peak calling using callpeak function from MACS3 (v3.0.2) with parameters: -- call-summits-g hs. Peaks overlapping regions in the hg38-blacklist.v2^61^ were removed from analyses.

### CUT&RUN

CUT&RUN samples and libraries were performed using the Epicypher Cut and Run kit (Epicypher, 14-1048-48rxn) and H3K9me3 primary antibody (Abcam, ab8898) according to manufacturer’s instructions. 500,000 cells were used per reaction and were permeabilized with 0.05% digitonin. IgG negative control antibody was provided in the kit. Libraries were sequenced on Illumina NextSeq platform with paired-end 75bp reads. The sequence data were trimmed by Trimmomatic (v0.40) to remove adaptor and then mapped to the hg38 assembly of the human genome using bowtie2 (v2.5.4). Duplicate reads were removed using picard-tools. Because the ecDNA amplicon region has substantially higher sequencing depth, which would reduce peak-calling *q*-values compared to chromosomal regions, we subsampled reads from the ecDNA amplicon according to ecDNA copy number to equalize sequencing depth between the ecDNA amplicon and chromosomal regions. The subsampled bam file was used for H3K9me3 peak calling using callpeak function from MACS3 (v3.0.2) with parameters:-g hs --keep-dup-q 0.01 --broad-cutoff 0.1 --min-length 1000 --max-gap 1000.

### ChIP-seq

ChIP-seq experiments were performed as previously described^10^. Primary antibodies used for chromatin immunoprecipitation during the ChIP-seq sample preparation: Lamin B1, Invitrogen, 702972; H3K9me3, Active Motif, 39065; H3K27ac, Abcam, ab4729; HP1a, Abcam, ab109028; HP1b, Abcam, ab270988; HP1γ, Sigma-Aldrich, 05-690. Sequencing libraries were prepared using the NEBNext Ultra II DNA Library Prep Kit (NEB, E7645) according to manufacturer’s instructions and sequenced on Illumina NovaSeq 6000 platform with paired-end 150 bp reads. The sequencing data were trimmed by TrimGalore (v0.6.10) to remove adaptor and then mapped to the hg38 assembly of the human genome using bwa (v0.7.17-r1188). Duplicate reads were removed using picard-tools. ChIP-seq signal was converted to the bigwig format using deepTools (v3.5.4.post1). ChIP-seq tracks were plotted using pyGenomeTracks^67^ (v3.9), and H3K9me3 peak heatmaps were plotted using deepTools. H3K9me3 peaks were called using callpeak function from MACS3 (v3.0.2) with parameters:-g hs --keep-dup-q 0.1 --min-length 1000 --max-gap 1000. To define High, Mid and Low H3K9me3 regions in **Figure 3H**, the whole genome was partitioned into 1-kb bins and H3K9me3 ChIP signal was quantified for each bin. Bins were then stratified based on their H3K9me3 levels: the top one-third of bins were classified as High H3K9me3 regions, the bottom one-third as Low H3K9me3 regions, and the remaining one-third as Mid H3K9me3 regions. LAD regions on chrDNA were inferred from Lamin B1 ChIP-seq, with LADs called in genomic bins where log2(treated/input) > 0 (**Figure S1C**).

### Immunofluorescence (IF) and fluorescence *in situ* hybridization (FISH)

Unless otherwise indicated, cells were grown on 12-mm glass coverslips coated with 10 µg/mL poly-D-lysine, fixed in 4% paraformaldehyde for 10 min at room temperature, and mounted using ProLong Gold antifade mountant following staining.

#### DNA FISH

For DNA FISH of cultured cells, fixed samples were permeabilized in 0.5% Triton X-100 in PBS for 15 min, and dehydrated through a graded ethanol series (70%, 85%, 100%). Labeled DNA probes targeting *MYC* and/or *FGFR2* (Empire Genomics) were diluted 1:10 in hybridization buffer and applied to samples. Cellular DNA was denatured at 80°C for 10 min followed by overnight hybridization at 37°C in a humidified chamber. After hybridization, samples were washed once in 0.4x SSC and twice in 2x SSCT (2x SSC supplemented with 0.1% Tween-20) to remove nonspecific binding. DNA FISH signals were detected using green-and red-labeled probes with nuclei counterstained with 10 µg/mL Hoechst. Samples were cured at room temperature for ∼24 h and stored at-20°C prior to imaging.

For DNA FISH of metastatic adenocarcinoma (omental nodule) patient tissue, the tumor was identified by a retrospective review of cases in the Stanford Pathology archive that had been identified as *MYC*-amplified on the Stanford Tumor Actionable Mutation Panel (STAMP). ecDNA status was determined empirically based on *MYC* DNA FISH signal appearance. DNA FISH was performed as previously described^52^ on 4 tissue cores of the tumor block. In brief, FFPE section was deparaffinized by two 5-min incubations in Histo-Clear (Electron Microscopy Sciences 64110), followed by 5 min in 100% ethanol and 5 min in 70% ethanol. The slide was then incubated in 0.2 N HCl for 20 min followed by a 20min incubation in 10 mM citric acid pH 6.0 in a vegetable steamer (90–95 °C). After brief wash in 2× SSC and treatment with proteinase K digestion buffer (1:100 dilution of proteinase K NEB P8107 in TE buffer) for 1 min at room temperature, the slide was then dehydrated by incubation for 2 min each in 70%, 85% and 100% ethanol. The *MYC* FISH probe (Empire Genomics MYC-20-GR) was diluted 1:5 in hybridization buffer (Empire Genomics), added to the slide, and covered with a coverslip. Denaturation was perfromed at 75 °C for 5 min followed by overnight hybridization at 37 °C in a humidified chamber. The slide was washed twice in 0.4× SSC + 0.3% Igepal630 (5 min, 40–60 °C) and then in 2× SCC + 0.1% Igepal630 (5 min, room temperature) followed by treatment with a TrueVIEW Autofluorescence Quenching kit (Vector laboratories SP-8400) according to the manufacturer’s directions for 2 min and then washed in 2× SSC (5 min, room temperature). The slide was mounted with ProLong Gold antifade with DAPI (ThermoFisher P36931). Slides were imaged on a Leica Thunder DMi8 microscope with z-step of 0.426 µm. Image merging, small volume computational clearing, and maximum intensity projections were performed using the LASX software. This component of the study was approved by the Stanford University Institutional Review Board (number 69198). DNA FISH in neuroblastoma patient tissue samples were performed as previously described^68^. DNA FISH in glioblastoma patient tissue samples were performed as previously described^69^.

#### RNA FISH

RNA FISH was performed following the Stellaris RNA FISH protocol, with minor modifications. Fixed samples were permeabilized in 70% ethanol for at least 1 h at 4°C. Samples were equilibrated in Wash Buffer A for 2-5 min and hybridized with custom-designed Stellaris RNA FISH probes targeting *MYC* intron 2 diluted 1:50 in hybridization buffer at 37°C overnight in a humidified chamber. The design of the RNA FISH probe has been described previously^3^. Following hybridization, samples were washed in Wash Buffer A at 37°C for 30 min, counterstained with 10 µg/mL Hoechst (in Wash Buffer A), cured at room temperature for ∼24 h, and stored at-20°C prior to imaging.

#### Immunofluorescence-DNA fluorescence in situ hybridization (IF-FISH)

For combined IF and DNA FISH, fixed samples were permeabilized in 0.5% Triton X-100 in PBS for 15min and blocked in 3% BSA in PBST (PBS supplemented with 0.1% Tween-20) for 1hr at room temperature. Samples were incubated with H3K9me3 primary antibody (1:500 dilution) in blocking buffer overnight at 4°C in a humidified chamber. After three washes in PBST (5 min each), samples were incubated with Alexa Fluor 647-conjugated secondary antibody (1:500 dilution) in blocking buffer for 30 min at room temperature. Following secondary antibody incubation, samples were washed with PBS and post-fixed in 4% paraformaldehyde for 20 min prior to DNA FISH processing. Samples were then permeabilized with ice-cold 0.7% Triton X-100 in 0.1M HCl (diluted in PBS) for 10 min on ice, followed by DNA denaturation in 1.9M HCl for 30min at room temperature. Samples were then dehydrated through a graded ethanol series (70%, 85%, 100%). The *MYC* DNA probe (Empire Genomics) was denatured at 75°C for 3 min, allowed to cool at room temperature for 5 min, and applied to samples for overnight hybridization at 37°C in a humidified chamber. After hybridization, samples were washed once in 0.4x SSC and twice in 2x SSCT (2x SSC supplemented with 0.1% Tween-20) to remove nonspecific binding. Nuclei were counterstained with 10 µg/mL Hoechst, and samples were cured at room temperature for ∼24 h and stored at-20°C prior to imaging.

### Microscopy

Unless otherwise noted, all images were collected on a Nikon Ti2 inverted microscope (Stanford University Cell Sciences Imaging Core Facility, RRID:SCR_017787) configured for spinning disk confocal imaging using a Plan Apo λ 60x oil-immersion objective (NA 1.4). Images were collected with a Photometrics Kinetix sCMOS camera controlled by NIS-Elements AR (v6.10.02). Z-stacks were acquired with 15-20 optical sections at 0.5 µm (PC3) or 1 µm (COLO320, SNU16) step size. Images were recorded at 16-bit depth with 1×1 binning at a spatial sampling of 0.108 µm per pixel.

### FISH imaging data analysis

#### Image processing

All imaging analyses were performed using custom-developed pipelines. DNA FISH Z-stacks were first maximum-intensity projected, and these max-projection images were used to compute per-channel flat-field correction images, which were applied to both max-projected and single-Z images.

#### Segmentation and background correction

Nuclei were segmented on flat-field corrected max-projection images using Cellpose with a custom-trained model. Segmented nuclei were filtered to remove objects with extreme size, irregular shape, or border contact. FISH signals were segmented using either custom developed segmentation pipeline or supervised pixel classification with custom-trained ilastik models. Signals outside of nuclei region were excluded from downstream analysis. Background intensity was estimated by random sampling of regions excluding nuclei and high-intensity signal, and all channels were background-corrected prior to quantification.

#### Single-Z Plane Extraction for Per-Nucleus Analysis

To enable single-plane analysis, an optimal focal Z-plane was determined for each nucleus by combining a field-of-view level Z-drift map with per-nucleus Z-intensity profiling. Only nuclei for which both methods identified similar focal plane were retained. Nuclear and FISH segmentation were repeated on these single-Z images using custom models.

#### Object-level Quantification and Spatial Metrics

Individual FISH objects were identified on single-Z images, and object-level metrics, including intensity, size, and distance to the nuclear boundary were quantified on a per-nucleus basis.

#### Radial Distribution Analysis

For each sample, the radial distribution of ecDNA was generated by aggregating the distances of individual ecDNA FISH-positive pixels from the nuclear edge across all nuclei, and calculating the normalized frequency of ecDNA FISH-positive pixels as a function of distance from the nuclear edge (**Figure 1C**). The random distribution was generated by assuming that, within each nucleus, all pixels had an equal probability of being ecDNA FISH-positive, thereby representing the expected radial distribution if ecDNA FISH signals were uniformly distributed throughout the nucleus. Confidence intervals of the distributions were estimated by nucleus-level resampling with replacement (95% percentile interval).

To assess the non-randomness of the radial distribution of FISH signals, a within-nucleus permutation test was performed. For each nucleus, the observed number of FISH-positive pixels was recorded, and FISH labels (that is, whether a pixel was FISH-positive) were randomly reassigned among nuclear pixels by sampling without replacement while preserving the per-nucleus FISH count. The randomly reassigned FISH-positive pixels were pooled across nuclei to generate null radial distance distributions. The 1D Wasserstein distance between the observed and permuted radial distance distributions was computed for each permutation, and permutation *P* values were obtained from the resulting null distribution (5000 permutations).

To compute the FISH signal enrichment curves (FISH/Random) shown in **Figure 1D**, the radial distribution profile of FISH signals was divided by the corresponding random distribution. Confidence intervals of the distributions were estimated by nucleus-level resampling with replacement (95% percentile interval). To compare the FISH signal enrichment curves between DM and HSR samples, the enrichment curves were first smoothed using a cubic spline and the midpoint of the region where enrichment exceeded 1 was defined as the enrichment peak center. The position difference of enrichment peak centers between DM and HSR samples were assessed using a nucleus-level permutation test (2000 permutations), in which nucleus labels (that is, whether the nucleus is from DM or HSR) were randomly reassigned while preserving sample sizes of DM and HSR to generate a null distribution, and the observed position difference in enrichment peak center was compared to the null distribution.

### CRISPR-DNA FISH optical screening

#### CRISPR perturbation and cell preparation

For each gene target, a pool of 3 sgRNAs (EditCo) was used to ensure efficient gene knockout. The sequences of sgRNAs are provided in **Table S1**. COLO320DM cells stably expressing mCherry-NLS were nucleofected with *in vitro*-assembled Cas9 ribonucleoprotein complexes containing arrayed CRISPR sgRNAs, while a GFP-NLS-expressing control population underwent mock nucleofection. Cells were allowed to recover and expand for four days post-nucleofection. Three biological replicates were performed per gene target, including a control condition consisting of mock-nucleofected mCherry-NLS and GFP-NLS cells.

#### Internal spike-in control and plate setup

To control for technical variability across wells, an internal spike-in strategy was employed. CRISPR-perturbed mCherry-labeled cells were mixed 1:1 with mock-nucleofected GFP-labeled control cells and co-seeded into the same wells of 96-well imaging plates four days post-nucleofection. After an additional 24-hour incubation, cells were fixed with 4% paraformaldehyde for 10 min and washed with PBS. This design ensured that control and perturbation conditions experienced identical experimental handling, hybridization conditions, and imaging parameters within each well.

#### 96-well DNA FISH

To enable high-throughput DNA FISH in a 96-well format, the standard DNA FISH protocol was modified to substantially reduce probe hybridization volume, the primary limitation preventing cost-effective implementation in multiwell formats (typically 10-15 µL per well in 384 well formats). DNA FISH was otherwise performed as described in the DNA FISH Methods section, with the following modifications for 96-well imaging plates: after probe application (3.5 µL per well), a 7-mm glass coverslip was placed over each well and sealed with rubber cement to prevent evaporation during high-temperature denaturation. Cellular DNA was denatured at 80°C for 13 min, followed by overnight hybridization at 37°C in a humidified chamber. This optimization preserved robust hybridization efficiency while enabling sufficient cell numbers per condition to capture ecDNA’s intrinsic heterogeneity.

#### Sequential imaging for cell identity assignment

Because the DNA denaturation step in DNA FISH abolishes fluorescent protein signals, a sequential imaging strategy was used to preserve cell identity. Fluorescent protein signals (GFP and mCherry) were imaged prior to DNA FISH to record cell identity and spatial coordinates using a Leica DMi8 widefield microscope controlled by Leica LAS X (AF60000LX), equipped with a Plan-Apochromat 10x/0.45 N.A. dry objective and a Leica DFC9000 GT sCMOS camera. Fluorescence illumination was provided by an LED light engine, and images were recorded as 16-bit widefield fluorescence images. DNA FISH was then performed as described above, followed by imaging of ecDNA signals as described in the Microscopy Methods section. Due to the utilization of two distinct microscopes, Hoechst channel is also collected separately with Leica DMi8 after FISH step to facilitate image registration.

#### Image analysis

All image analysis was performed using custom-developed Python pipelines. Pre-FISH images containing fluorescent protein signals (GFP and mCherry, collected with Leica DMi8) and post-FISH images containing DNA FISH signals (collected with Nikon Ti2) were computationally registered to assign CRISPR perturbation identity to individual cells, assisted by an additional post-FISH Hoechst dataset (collected with Leica DMi8) to enable accurate cross-platform alignment. Identity assignment was restricted to spatially matched nuclear pairs within a defined distance threshold. DNA FISH radial distribution analysis was performed as described in the *FISH Imaging Data Analysis* section. For each perturbation, radial enrichment profiles were computed separately for control (GFP-labeled) and CRISPR-perturbed (mCherry-labeled) cells. Only datasets containing at least 50 cells per group were retained for downstream comparison to ensure robust estimation of population-level effects. Screen-level comparisons were summarized by calculating the log2 fold change in radial enrichment metrics between perturbed and control populations. These values were aggregated across biological replicates and visualized as heatmaps to enable comparative analysis across gene perturbations.

### Simulation of nuclei containing ecDNA-like foci

To generate synthetic nuclei with ecDNA-like foci exhibiting controlled spatial organization, we performed pixel-level simulations in which both radial enrichment affinity and intermolecular interaction strength were systematically varied. All fixed simulation parameters, including nuclear size, ecDNA copy number per nucleus, spot size, and minimum inter-spot distance, were estimated from experimental measurements in COLO320DM cells to match the spatial scale and density observed in this system.

#### Peripheral attraction modeling

Nuclei were modeled as circular domains of fixed radius and discretized on a square pixel grid. Radial enrichment was specified using a family of spline-based enrichment profiles derived from the experimentally measured ecDNA radial enrichment curve in COLO320DM cells (**Figure 1D**). These profiles preserved the overall shape of the experimental distribution while varying its amplitude, thereby representing graded peripheral enrichment affinities. For each simulation, ecDNA seed locations were sampled probabilistically across nuclear pixels using sampling weights derived from the spline-based enrichment profile as a function of normalized radial position. A fixed number of ecDNA seeds were placed per nucleus, subject to a minimum inter-spot distance constraint.

#### Intermolecular interaction modeling

To model intermolecular interactions between ecDNA foci, a short-range, distance-dependent attraction term was applied during ecDNA seed placement. For each candidate location, the sampling probability was biased toward proximity to previously placed ecDNA foci within a defined interaction radius. The intermolecular interaction strength parameter scaled the magnitude of this attraction, increasing the tendency for newly placed ecDNA foci to be positioned closer to existing ones without altering the prescribed radial distribution.

#### Simulation and image analysis

Large synthetic datasets were generated by simulating 10,000 nuclei per condition, which were assembled into tiled images for downstream image analysis using the same pipeline as experimental FISH data, except that flat-field correction, segmentation, and background correction were not required.

### Pore-C

#### Cell preparation

For **Figure S7C**, two million cells were fixed in 1% PFA for 10 min at room temperature, and quenched by adding glycine to the sample suspension for a final concentration of 125mM glycine. Cells were incubated at room temperature for 5 minutes, then chilled on ice for a further 10 minutes with regular, gentle agitation. The cells were washed 3 times with cold PBS and pelleted by centrifugation at 500 g at 4°C for 5 minutes. For **Figure 5B**, monoclonal COLO320DM-FUCCI cells were incubated with 50µM BrdU in culture medium for 2 h at 37 °C prior to fixation, and cells were washed once with 1X PBS, trypsinized, and the cell mixture was passed through a 40µm cell strainer into a 50mL tube. The cells were spun down and washed once with 1X PBS, and resuspended in 1X PBS containing 0.1% FBS and LIVE/DEAD™ Fixable Far Red Dead Cell Stain Kit, for 633 or 635 nm excitation (Invitrogen, #L34973). The cells were incubated at room temperature for 20 mins in the dark to label dead cells. Cells were then washed once with 1X PBS and fixed with freshly prepared 1% paraformaldehyde for 10 mins at room temperature. The reaction was quenched by the addition of glycine at a final concentration of 125mM for 5 mins at room temperature. The cells were then incubated on ice for another 10 mins with gentle agitation, spun down and washed thrice with cold 1X PBS. Cells were resuspended in Hanks’ Balanced Salt Solution (Gibco, 14025092) for fluorescence-activated cell sorting to obtain 1M cells in early and late S phase respectively, as indicated by the FUCCI indicator. Cell sorting for this project was done on instruments in the Stanford Shared FACS Facility (RRID: SCR_017788). Sorted early-S and late-S cells were pelleted by centrifugation at 500 g at 4°C for 5 minutes.

#### Pore-C

Pore-C was performed as previously described^38^. Fixed and pelleted cells were resuspended in DW buffer (0.02% digitonin, 20 mM HEPES-potassium hydroxide buffer, pH 7.5, 150 mM sodium chloride, 0.5 mM spermidine, 1 Roche cOmplete EDTA-free tablet (Roche, 11873580001) per 50 ml buffer and 0.1% BSA) and incubated on ice for 5 min. Cells were centrifuged for 3 minutes at 500 g at 4°C, and washed once in TW buffer (0.1% Tween-20, 20 mM HEPES-potassium hydroxide, pH 7.5, 150 mM sodium chloride, 0.5 mM spermidine, 1 Roche cOmplete EDTA-free tablet per 50 ml buffer and 0.1% BSA), then spun down for 3 minutes at 500 g at 4°C. The cell pellet was washed again in 1.5X digestion reaction buffer (1.5X CutSmart buffer for NlaIII or 1.5X NEBuffer DpnII for DpnII), spun down for 3 minutes at 500 g at 4°C, and resuspended in 300 μL of 1.5X digestion reaction buffer. To denature the chromatin, 33.5 μl of 1% (wt/vol) SDS (Thermo Fisher Scientific, 15553027) was added and incubated for 10 min at 65 °C and placed on ice immediately afterward. Next, 37.5 μl of 10% (vol/vol)

Triton X-100 (Sigma-Aldrich, 93443) was added and incubated for 10 minutes on ice. One U /μl of DpnII (NEB, R0543M, **Figure 5B**) or NlaIII (NEB, R0125L, **Figure S7C**) was added to the cells with nuclease-free water to achieve a final 1× digestion reaction buffer in 450 μl, and cells were incubated in at 37 °C for 18 h with periodic ∼ 500 r.p.m. rotation (15 s every 15 min) to prevent condensation inside the lid. The restriction digests were heat inactivated at 65 °C for 20 min. Digested cells were spun down for 3 minutes at 300 g, and supernatant was carefully removed. Proximity ligation was performed by resuspending the sample in ligation reaction (1X T4 DNA Ligase Reaction Buffer (NEB, M0202L), 0.1 μg/μl BSA, 20 U/μl T4 DNA ligase (NEB, M0202L)) and incubated at 16 °C for 4 h. The sample was spun down for 3 minutes at 300 g, and supernatant was carefully removed. The sample was then resuspended in reverse-crosslinking buffer (5% (vol/vol) Tween 20, 0.5% (wt/vol) SDS, 1 μg/μl Proteinase K (NEB, P8107S), 0.5X PBS), and incubated in a 56 °C incubator for 18 h. DNA was subsequently purified by phenol–chloroform extraction followed by ethanol precipitation. Nanopore sequencing libraries were prepared using the Native Barcoding Kit 24 V14 (Oxford Nanopore Technologies, SQK-NBD114.24) following the manufacturer’s instructions with the exception that during the FFPE repair and Ultra II DNA repair and end-prep step, the reaction was incubated at 20 °C for 15 min instead of 5 min. Libraries were sequenced on an Oxford Nanopore PromethION 24 platform.

### Pore-C data analysis

#### Basecalling and alignment

Base calling of canonical bases was performed using Dorado (v0.9.1) and models dna_r10.4.1_e8.2_400bps_sup@v5.0.0. Base called reads were aligned to hg38 reference genome using minimap2 (v2.26-r1175). BrdU incorporation was detected using DNAscent^39^ (v4.0.3).

#### Classification of nuclear-peripheral ecDNA reads and nuclear-interior ecDNA reads

Unlike Hi-C which captures pair-wise contacts, Pore-C captures multi-way contacts between DNA fragments, providing proximity information of each DNA fragment. For example, as shown in **Figure S6A**, DNA Fragment 1, Fragment 2, Fragment 3, and Fragment 4 are spatially in close proximity within the nucleus. In Pore-C, they are ligated together as a single long DNA molecule for nanopore long-read sequencing (**Figure S6A**). After sequence alignment, each fragment (e.g., Fragment 1) is referred to as a “read”, while other fragments on the same sequenced DNA molecule (e.g., Fragment 2, Fragment 3 and Fragment 4) are called “contact loci” of the read (i.e., Fragment 2, Fragment 3 and Fragment 4 are the contact loci of Fragment 1). The contact loci of each read are summarized in the ‘SA’ tag in the aligned bam output of minimap2.

The spatial location of a DNA fragment in the cell can be inferred from its contact loci. **Figure S6B** illustrates how Pore-C reads aligned to the ecDNA amplicon region are classified as nuclear-peripheral or nuclear-interior ecDNA reads. EcDNA read A has 10 chromosomal contact loci. Notably, 10 out of the 10 chromosomal contact loci are located within constitutive lamina-associated domain (cLAD) regions. When compared to all reads, ecDNA read A shows significantly enriched contacts with cLAD regions (hypergeometric test *p* < 0.01), suggesting that ecDNA read A originates from an ecDNA molecule surrounded by cLAD chromatin at nuclear periphery (**Figure 5C**). Therefore, ecDNA read A is classified as a nuclear-peripheral ecDNA read. Similarly, ecDNA read B exhibits significantly enriched contacts with constitutive inter-lamina-associated domain (ciLAD) regions (hypergeometric test *p* < 0.01), indicating that ecDNA read B originates from an ecDNA molecule in proximity to ciLAD chromatin at the nuclear interior. Therefore, ecDNA read B is classified as a nuclear-interior ecDNA read. Previously annotated cLAD and ciLAD regions were obtained from the previous study^20^ and lifting over to hg38 using the liftOver tool (UCSC).

### Quantification of DNA replication timing and ecDNA-ecDNA contact frequency using Pore-C

For each read, DNAscent^39^ (v4.0.3) reported the probability that each thymidine corresponded to BrdU incorporation. A read was classified as BrdU+ if the 70^th^ percentile of BrdU probability across all thymidines in that read exceeded 0.5. Replication timing was quantified as the ratio of the fraction of BrdU+ reads in late S-phase cells to the fraction of BrdU+ reads in early S-phase cells (**Figure 5D**). Higher replication timing ratios indicate later replication timing. For instance, the ratio greater than 1 indicates more DNA replication during late S phase than early S phase, consistent with late replication timing, whereas a ratio less than 1 indicates preferential DNA replication during early S phase.

To calculate ecDNA-ecDNA contact frequency using Pore-C data (**Figure S7C**), reads aligned to ecDNA amplicon were first classified as nuclear-peripheral and nuclear-interior ecDNA reads as above described. For each class (peripheral or interior), ecDNA-ecDNA contact frequency was calculated as the total number of contact loci falling within ecDNA amplicon regions across the corresponding reads, divided by the total number of the corresponding reads.

### Hi-C

COLO320DM Hi-C raw data was obtained from the previous publication^57^. Raw Hi-C sequencing data were processed using HiC-Pro (v3.1.0) pipeline to generate genome-wide contact matrices at multiple resolutions. Briefly, paired-end reads were aligned to the hg38 genome, filtered for valid interaction pairs, and deduplicated using the default settings of the HiC-Pro pipeline. The pipeline was configured to assign reads to DpnII restriction fragments and merge valid interaction pairs into one.allValidPairs file, and then build raw contact matrices by binning interaction counts into multiple resolutions of genomic bins. The Hi-C contact counts in **Figure 6D** were then quantified by summing interaction counts between ecDNA amplicon regions (ec-ec) or between ecDNA amplicon and LAD chrDNA regions (ec-LAD) using the 1-kb resolution raw contact matrix.

### RNA-seq

One to two million cells cultured in 6-well plates were directly lysed using Buffer RLT Plus (Qiagen, 74134), and total RNA was isolated using RNeasy Plus Mini Kit (Qiagen, 74134) following the manufacturer’s instructions. RNA-seq libraries were prepared using NEBNext Ultra II Directional RNA Library Prep Kit for Illumina (NEB, E7760S) with rRNA depletion using NEBNext rRNA Depletion Kit v2 (Human/Mouse/Rat) with RNA Sample Purification Beads (NEB, E7405L) according to the manufacturer’s protocol, and sequenced on Illumina NovaSeq 6000 platform with paired-end 150 bp reads. Sequencing reads were aligned to the human reference genome hg38 using STAR (v2.7.10b). Read counts aligned to each gene (gene body) were quantified using featureCounts (v2.1.1). Differential gene expression was calculated using PyDESeq2^70^ (v0.5.2) with a baseMean cutoff 100 to filter out very lowly expressed genes. The gene set enrichment analysis of ecGenes was performed using GSEApy^71^ (v1.1.11), with the Wald statistic for individual genes calculated by PyDESeq2 as the ranking metric.

### Nanopore WGS

One million WT or *LBR*-KO COLO320DM cells were prepared as described in the optical screening section. At 5 days after nucleofection, cells were harvested, and genomic DNA was purified using the MagAttract HMW DNA Kit (Qiagen, 67563) according to the manufacturer’s protocol. Nanopore sequencing libraries were prepared using the Native Barcoding Kit 24 V14 (Oxford Nanopore Technologies, SQK-NBD114.24) following the manufacturer’s instructions and sequenced on an Oxford Nanopore PromethION 24 platform. Basecalling of canonical bases was performed using Dorado (v0.9.1) with the dna_r10.4.1_e8.2_400bps_sup@v5.0.0 model. Basecalled reads were aligned to the hg38 reference genome using minimap2 (v2.26-r1175). EcDNA copy number was estimated as two times the ratio of the mean sequencing depth across ecDNA amplicon regions to the mean sequencing depth in non-amplicon genomic regions excluding the hg38-blacklist.v2^61^ regions.

### TCGA RNA expression analysis in ecDNA+ and ecDNA− samples

Amplicon status for TCGA samples was obtained from Amplicon Repository, based on AmpliconArchitect (AA) outputs (https://ampliconrepository.org/project/655bddb5bba7c92509525039). RNA-seq data were downloaded as TOIL-processed RSEM TPM values from the UCSC Xena Toil RNA-seq Recompute (https://toil-xena-hub.s3.us-east-1.amazonaws.com/download/tcga_RSEM_gene_tpm.gz). AA-derived amplicon classifications were integrated with RNA-seq data to stratify samples into ecDNA(+) and ecDNA(−) groups. Gene expression differences between groups were assessed for selected genes, and statistical significance was determined using the two-sided Mann–Whitney U test.

### Quantification and statistical analysis

Details of exact statistical analysis, tests, and other information can be found in the main text, figure legends, and Methods.

### Declaration of generative AI and AI-assisted technologies in the writing process

During the preparation of this work the authors used OpenAI ChatGPT in order to improve the readability and language of the manuscript. After using this tool/service, the authors reviewed and edited the content as needed and take full responsibility for the content of the published article.

---

**Figure S1.**
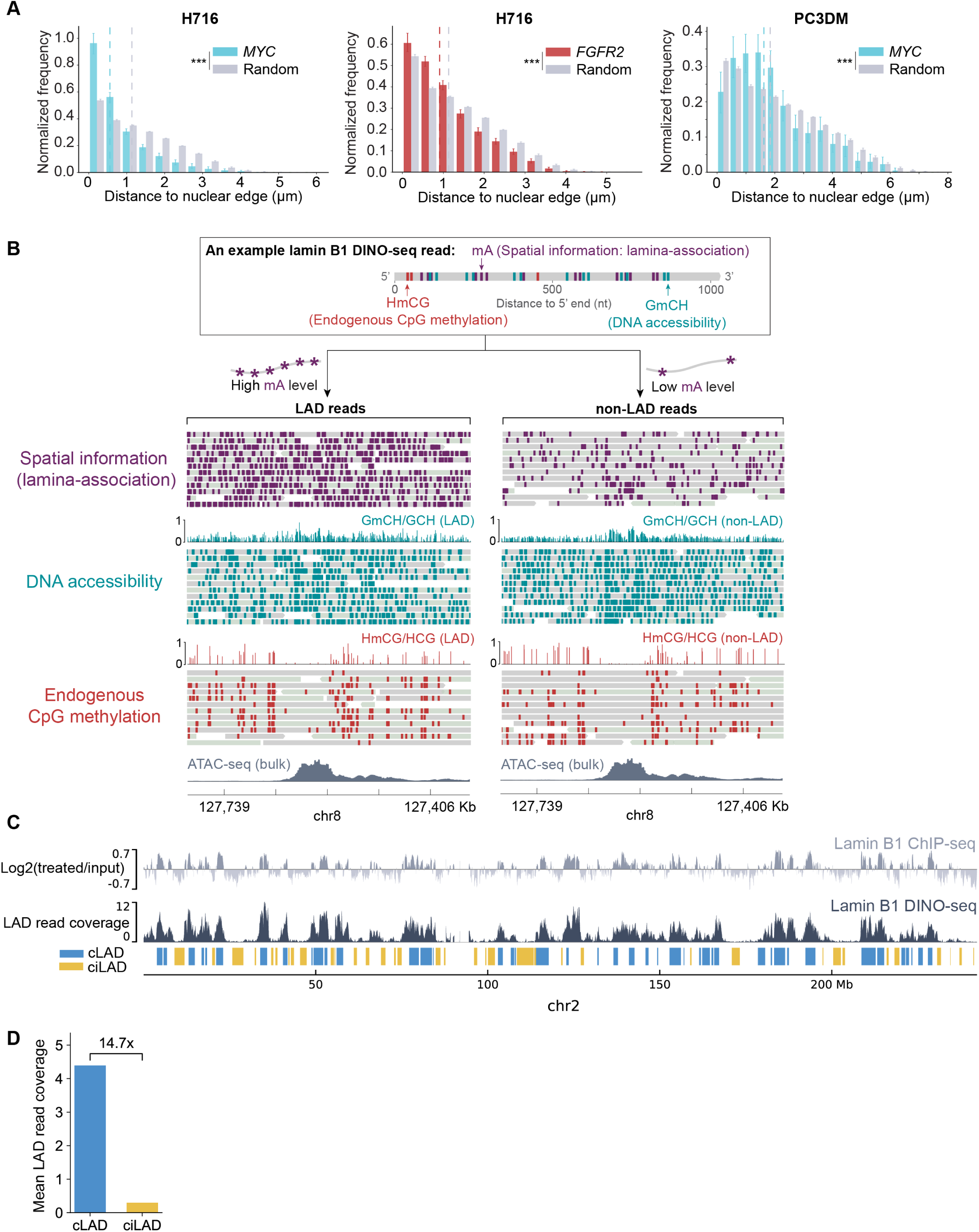
Quantification of DNA FISH visualization of ecDNA and example reads of lamin B1 DINO-seq, related to Figure 1 and Figure 2 (A) Radial distribution of ecDNA distances to the nuclear edge, quantified from the DNA FISH data from H716 and PC3DM cells in Figure 1B, and compared with a random distribution in which ecDNA FISH signals were uniformly distributed across the nucleus. Dashed lines indicate the median of each distribution. Error bars represent 95% confidence intervals. *** represents *p* < 0.001. *P* values were calculated using a two-sided permutation test (5000 permutations) comparing the observed and random radial distributions. H716, colorectal cancer; and PC3DM, prostate cancer. (B) Example reads from lamin B1 DINO-seq of COLO320DM cells. Each read was classified as a LAD read or non-LAD read based on its adenine methylation (mA) level. After the read classification, DNA accessibility and endogenous CpG methylation of LAD and non-LAD reads can be separately evaluated. Base modification mA (dark purple), GmCH (turquoise), HmCG (red) are labeled on each LAD and non-LAD reads. Each read is displayed with a grey or light-green background, indicating alignment to the plus or minus strand of the hg38 reference genome, respectively. Pseudo-bulk GmCH/GCH and HmCG/HCG tracks are plotted above the reads. Figure S2A explains how the pseudo-bulk tracks were calculated. (C) Comparison between lamin B1 ChIP-seq signals and LAD read coverage of lamin B1 DINO-seq across chromosome 2 in COLO320DM cells. Previously annotated constitutive LAD (cLAD) and constitutive inter-LAD (ciLAD) regions^20^ are indicated by blue and yellow boxes, respectively. (D) Comparison of mean LAD read coverage of lamin B1 DINO-seq in COLO320DM across previously annotated cLAD and ciLAD regions^20^ (excluding ecDNA amplicons and blacklisted regions^61^), which shows LAD reads are enriched at cLAD regions and depleted at ciLAD regions.

**Figure S2.**
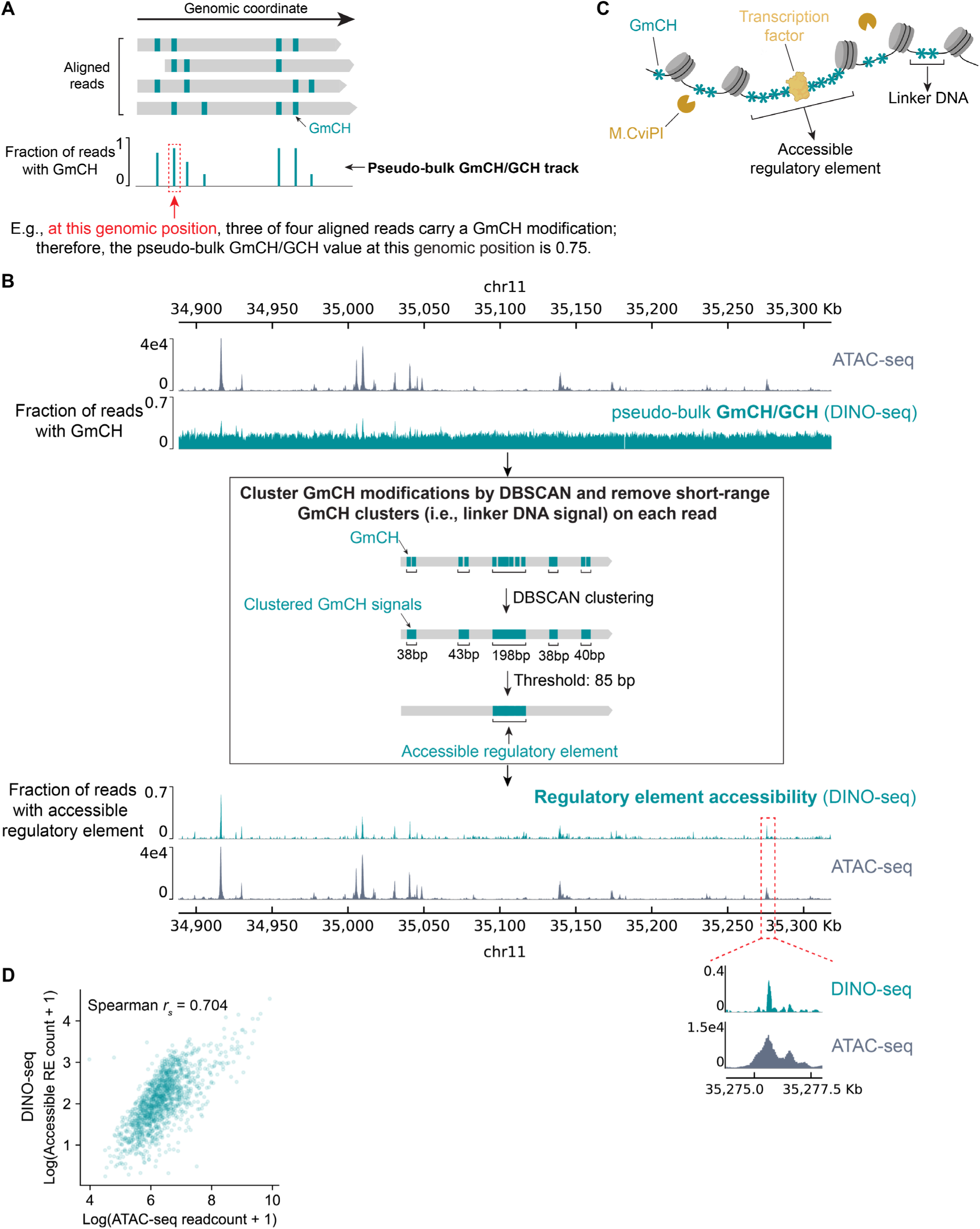
Nuclear-peripheral ecDNA accessibility analysis using lamin B1 DINO-seq, related to Figure 2 (A) Schematic of computing the pseudo-bulk GmCH/GCH track. At each GCH motif position across the genome, the fraction of reads that carry a GmCH modification at the position was calculated. This fraction is recorded as the pseudo-bulk GmCH/GCH value for the genomic position. (B) ATAC-seq, pseudo-bulk GmCH/GCH track and pseudo-bulk regulatory element accessibility track of lamin B1 DINO-seq from SNU16m1 cells, and schematic of filtering out short-range GmCH clusters corresponding to linker DNA for each read. To remove short-range GmCH clusters arising from linker DNA, the Density-Based Spatial Clustering of Applications with Noise (DBSCAN) was applied to identify discrete GmCH signal clusters along each read. Each cluster corresponds either to linker DNA between adjacent nucleosomes or to an accessible regulatory element. To distinguish these, clusters shorter than 85 bp, which represent linker DNA, were removed. The remaining GmCH clusters therefore reflect accessible regulatory elements. The pseudo-bulk regulatory element accessibility track was computed by calculating, at each genomic position, the fraction of reads carrying an accessible regulatory element overlapping with the position. The ATAC-seq track is shown twice for comparison with the pseudo-bulk GmCH/GCH track and the pseudo-bulk regulatory element accessibility track. The red dashed rectangle denotes a zoomed-in region comparing the pseudo-bulk regulatory element accessibility track from DINO-seq with the ATAC-seq track, illustrating the sharper peak profiles obtained by DINO-seq as a result of its near single-base-pair resolution. (C) Schematic illustrating that both accessible regulatory elements and linker DNA connecting adjacent nucleosomes are accessible to the GpC methyltransferase M.CviPI, and are consequently modified by the enzyme. (D) Correlation between ATAC-seq signal and lamin B1 DINO-seq accessible regulatory elements from SNU16m1 cells. Spearman correlation coefficient *r_s_* was computed using the counts of accessible regulatory elements (RE) and ATAC-seq reads in each 2-kb bin across ecDNA amplicon.

**Figure S3.**
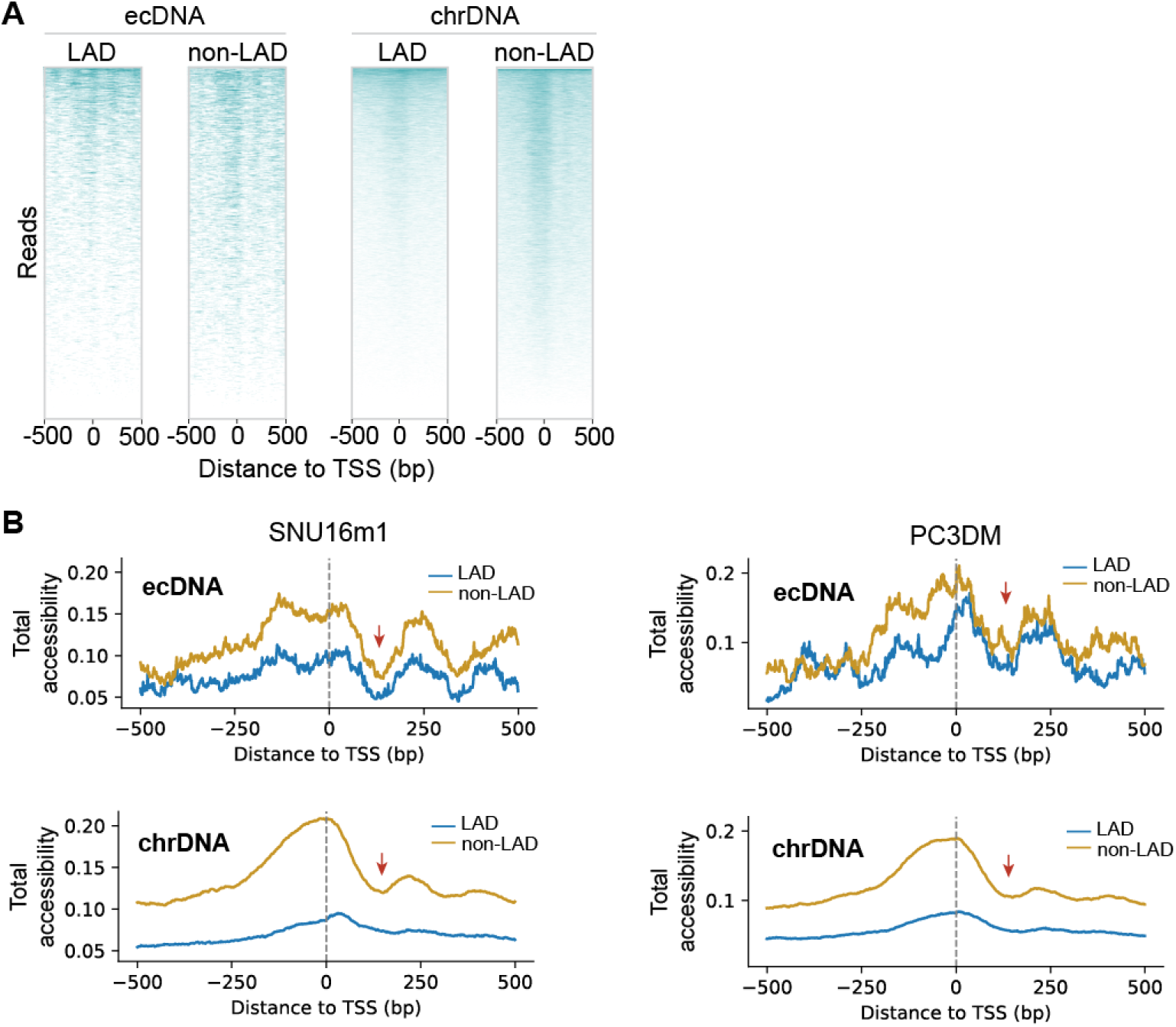
DNA accessibility analysis of lamin B1 DINO-seq reads at TSS, related to Figure 2 (A) Lamin B1 DINO-seq reads centered at TSS and sorted based on their accessibility at the TSS in COLO320DM cells. Turquoise color represents GmCH clusters from DBSCAN (including both accessible regulatory elements and linker DNA; see Figure S2B). (B) DNA accessibility analysis of lamin B1 DINO-seq reads at TSS in SNU16m1 and PC3DM cells. Line profiles show the total accessibility (i.e., the sum of regulatory element accessibility and linker DNA accessibility) of lamin B1 DINO-seq reads at each position around TSS. Red arrows indicate the +1 nucleosome footprints.

**Figure S4.**
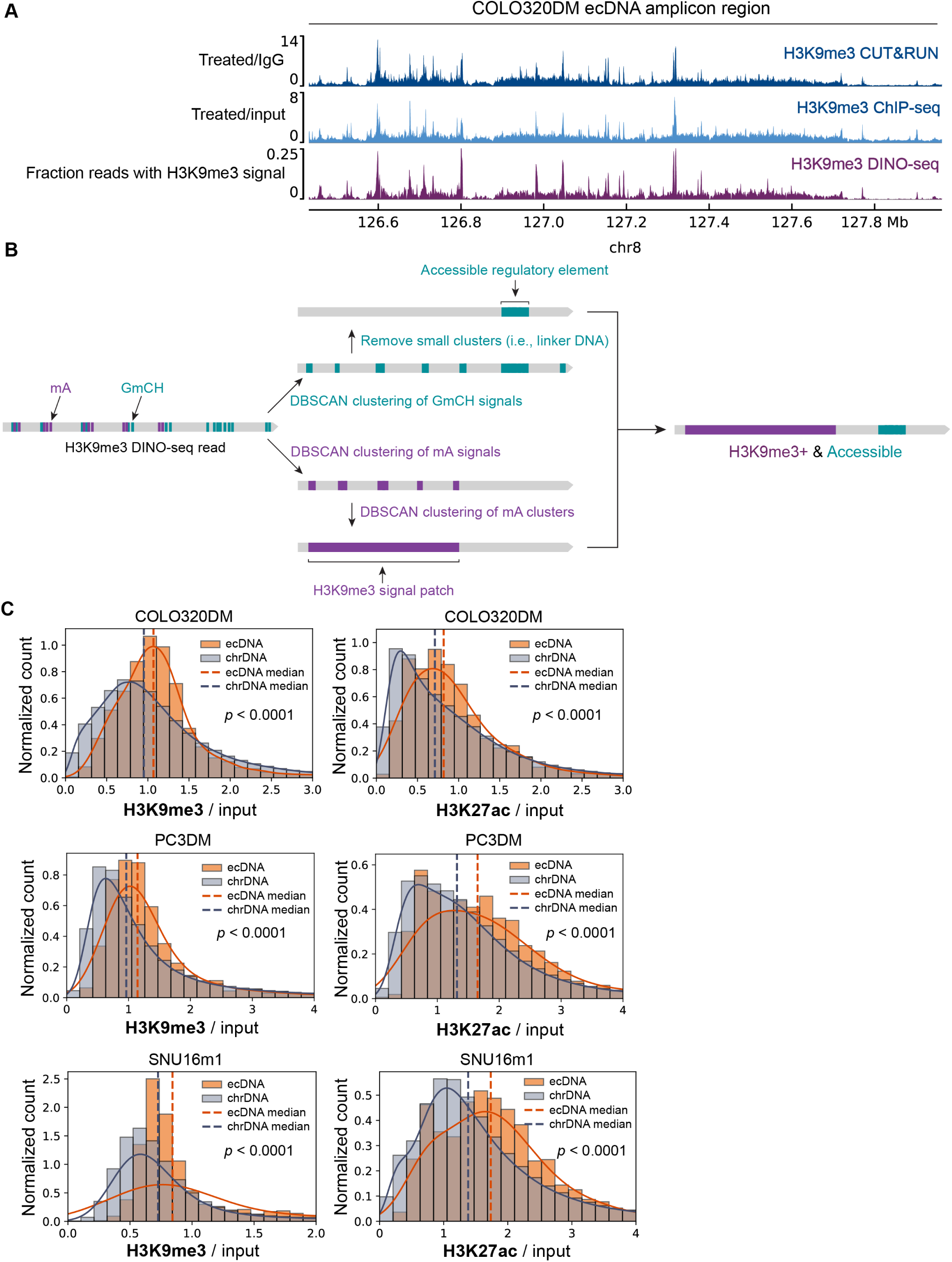
EcDNA histone modification landscape, related to Figure 3 (A) Comparison of H3K9me3 CUT&RUN, H3K9me3 ChIP-seq, and pseudo-bulk H3K9me3 signals of DINO-seq at the ecDNA amplicon region of COLO320DM cells. The pseudo-bulk H3K9me3 track was generated by computing, at each genomic position, the fraction of reads that have an H3K9me3 signal patch (identified as described in Figure S4B) overlapping with the position. (B) Schematic of H3K9me3 DINO-seq read-processing workflow for identifying accessible regulatory elements and H3K9me3 signal patch on individual reads. The GmCH and mA modifications on each read were analyzed separately. GmCH modifications were analyzed as described in Figure S2B to identify accessible regulatory elements. For mA modifications, two rounds of DBSCAN clustering were performed because H3K9me3 domains typically span broad genomic regions. In the first round of DBSCAN, individual mA events were grouped into discrete mA clusters. In the second round of DBSCAN, adjacent mA clusters were merged into extended H3K9me3 signals. See Methods for details. (C) Comparison of H3K9me3 and H3K27ac ChIP-seq signals on ecDNA and chrDNA regions in different cell lines. For this analysis, the whole genome was partitioned into 1-kb bins. H3K27ac and H3K9me3 ChIP-seq signals (i.e., treated/input) were quantified for each bin. The histograms show the distribution of H3K27ac or H3K9me3 ChIP-seq signal of ecDNA bins (i.e., 1-kb bins located within ecDNA amplicon regions) and chrDNA bins (i.e., 1-kb bins outside ecDNA amplicon regions). Dashed lines represent the median values of corresponding histograms. Solid curves represent kernel density estimates (KDE) of the signal distributions. *P* values were computed using two-sided Mann–Whitney U tests.

**Figure S5.**
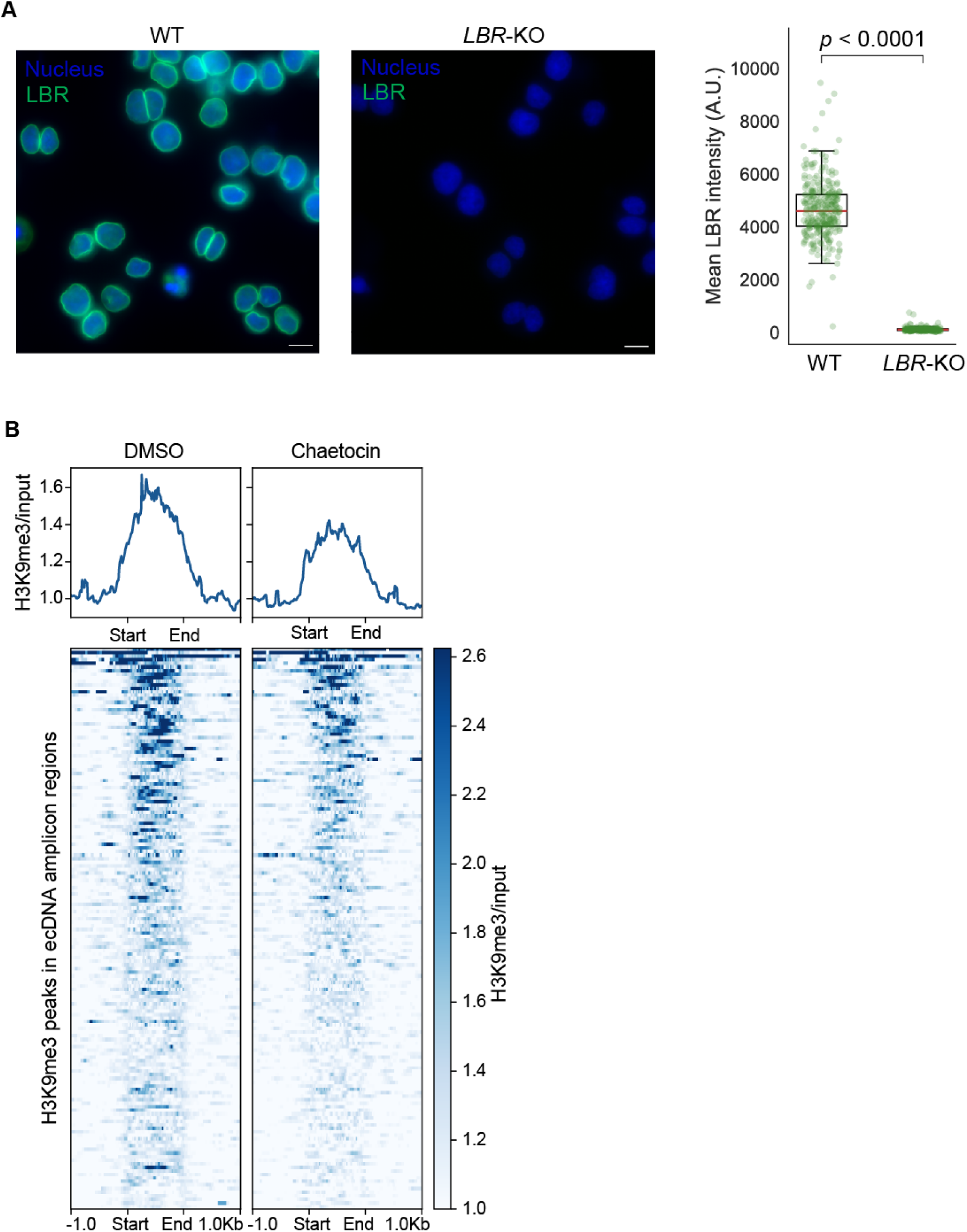
LBR immunofluorescence validates efficient knockout, and chaetocin treatment reduces H3K9me3 levels on ecDNA, related to Figure 4 (A) Representative images of LBR immunofluorescence (IF) in WT and *LBR*-KO cells, and quantification of LBR IF intensity confirming efficient depletion of LBR at 5 days after CRISPR knockout. In the box plots, each dot represents one cell. Box plots show the median and interquartile range (IQR), with whiskers extending to 1.5 × IQR. *P* value was calculated using a two-sided Mann– Whitney U test. (B) Heatmap analysis of individual H3K9me3 ChIP-seq peaks at ecDNA amplicon regions in COLO320DM cells treated with DMSO or chaetocin (Figure 4D). Aggregated H3K9me3/input signal profiles across these peaks are shown above the heatmaps.

**Figure S6.**
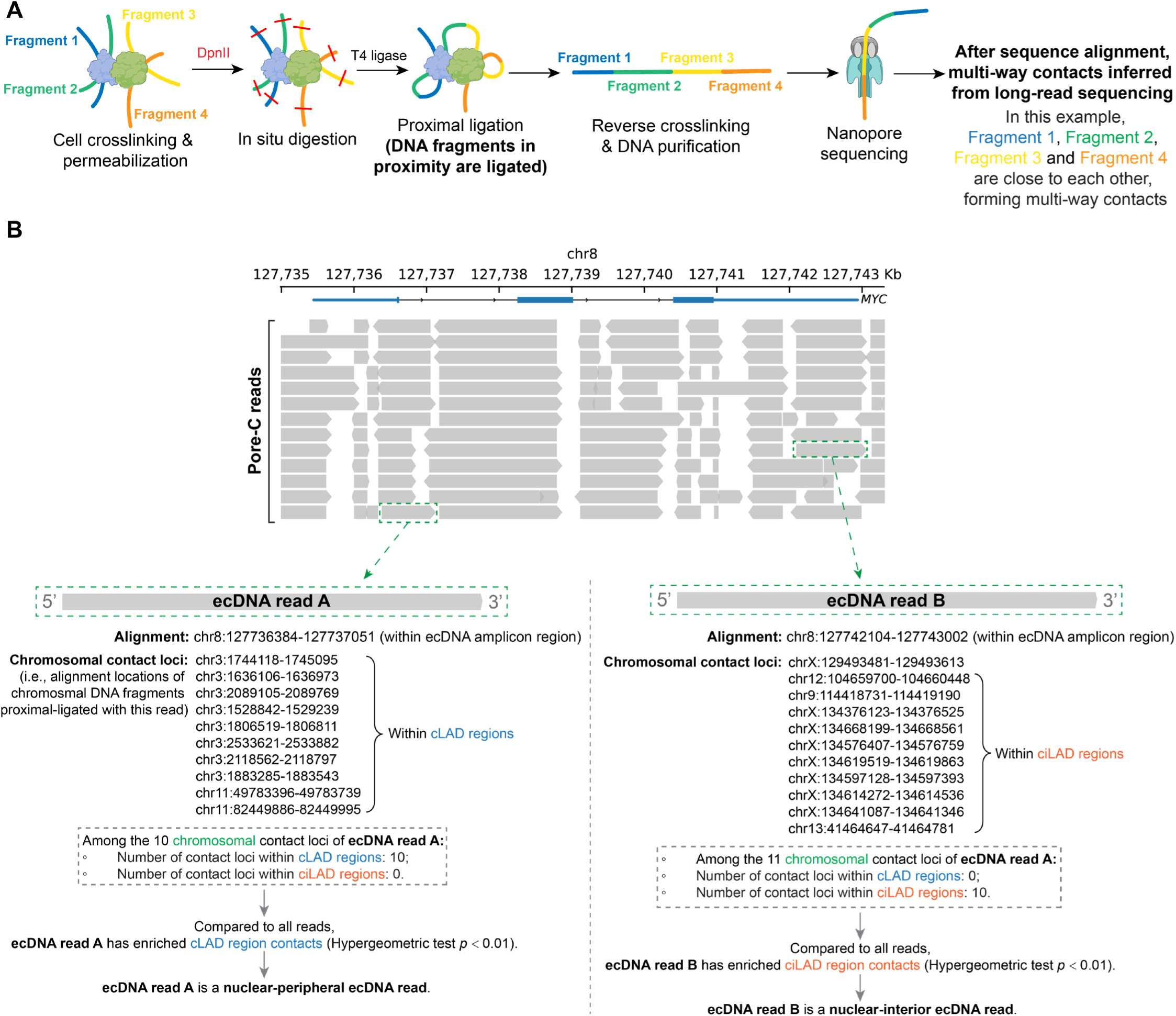
Pore-C read analysis, related to Figure 5 (A) Pore-C workflow. Cells are fixed by crosslinking and permeabilized, followed by a restriction enzyme (e.g., DpnII) treatment to digest genomic DNA. Next, the digested DNA fragments are proximally ligated. The DNA is then reversed crosslinked and purified for Nanopore sequencing. Unlike Hi-C, which captures pairwise chromatin contacts, Pore-C captures multi-way contacts among multiple DNA fragments, thereby providing higher-order proximity information^38^. For example, as shown in this schematic, DNA Fragment 1, Fragment 2, Fragment 3 and Fragment 4 are spatially proximal within the nucleus and become ligated into a single long DNA molecule during the Pore-C protocol. Nanopore long-read sequencing of this molecule yields a read composed of multiple aligned fragments. After alignment, each fragment (for example, Fragment 1) is considered a **read**, and the other fragments co-ligated on the same molecule (Fragments 2–4) are defined as **contact loci** of the read (i.e., Fragment 2, Fragment 3 and Fragment 4 are the contact loci of Fragment 1). (B) Example Pore-C reads aligned to the ecDNA amplicon region and their classification as nuclear-peripheral or nuclear-interior ecDNA. The spatial position of a DNA fragment can be inferred from the genomic locations of its contact loci. In the example shown, ecDNA read A contains 10 chromosomal contact loci, all of them fall within cLAD regions. Relative to the contact distribution of all reads, read A shows significantly enriched contacts with cLAD regions (Hypergeometric test *p* < 0.01), indicating that the ecDNA molecule is surrounded by cLAD chromatin at the nuclear periphery (Figure 5C). Thus, ecDNA read A is classified as a nuclear-peripheral ecDNA read. Conversely, ecDNA read B exhibits significantly enriched contacts with ciLAD regions (Hypergeometric test *p* < 0.01), suggesting that the ecDNA molecule resides near ciLAD chromatin in the nuclear interior. Therefore, ecDNA read B is classified as a nuclear-interior ecDNA read.

**Figure S7.**
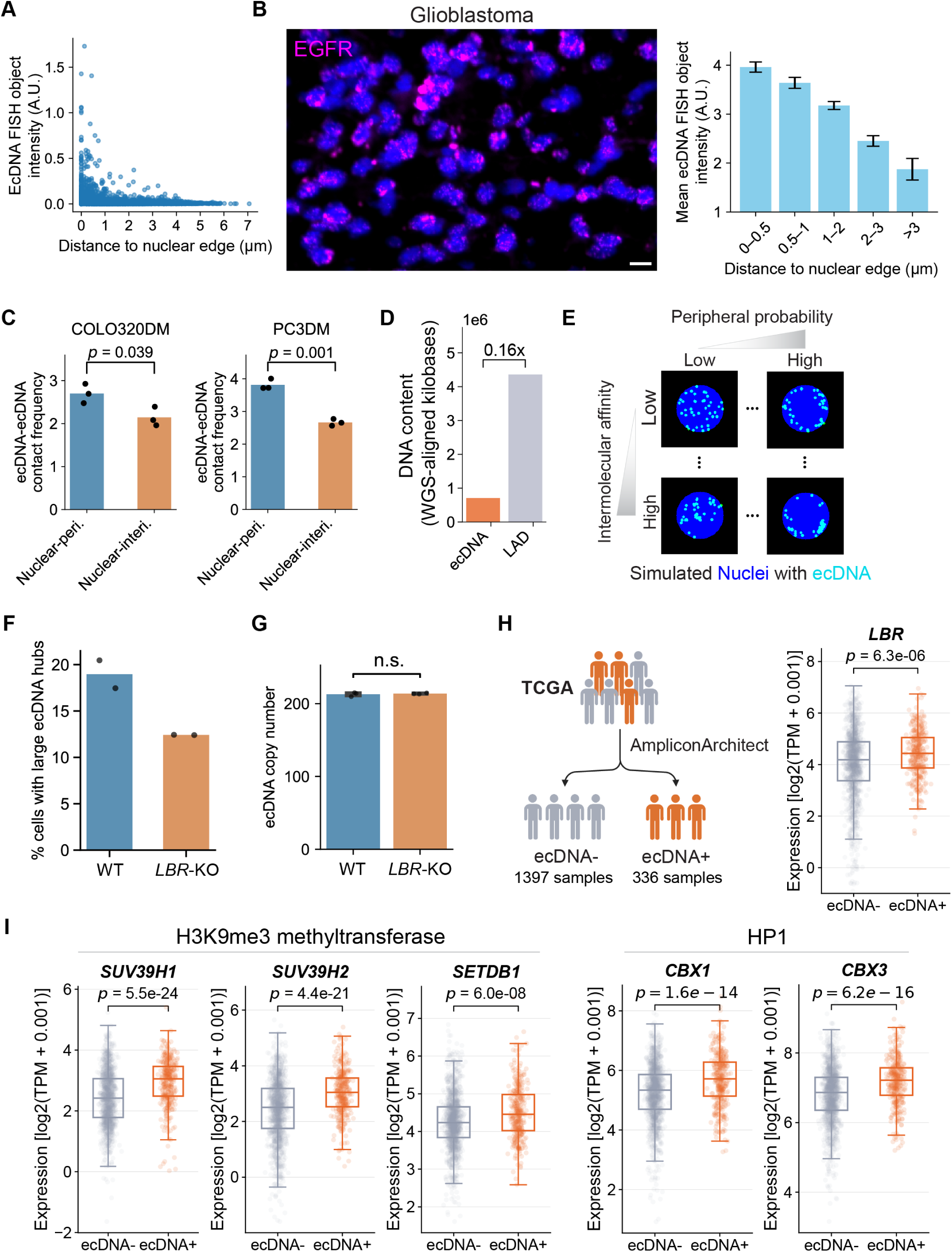
Nuclear-peripheral localization facilitates large ecDNA hub formation, and gene expression analysis of TCGA samples, related to Figure 6 (A) Scatter plot of total FISH intensity and distance to the nuclear edge of individual ecDNA FISH objects measured from DNA FISH in COLO320DM cells. Each dot represents an ecDNA FISH object. (B) Representative DNA FISH image from a glioblastoma patient, and bar plot of mean FISH intensity of individual ecDNA FISH objects as a function of distance to the nuclear edge. Error bars indicate standard error of the mean. (C) Pore-C analysis comparing ecDNA-ecDNA contact frequencies between nuclear-peripheral ecDNA and nuclear-interior ecDNA. Black dots represent independent replicates (n=3). *P* values were calculated using two-sided Student’s t-test (n=3). (D) Comparison of the difference in cellular DNA content between ecDNA and LAD chrDNA. Nanopore whole-genome sequencing (WGS) was performed, and the DNA content for ecDNA and LAD chrDNA was quantified as the total number of sequenced bases aligned to ecDNA amplicon regions or LAD chrDNA regions, respectively. LAD chrDNA regions were inferred from Lamin B1 ChIP-seq, with LADs called in genomic bins where log2(treated/input) > 0 (Figure S1C). The data indicate that LAD chrDNA contributes 6.25-fold more to the total cellular DNA content than ecDNA. (E) Representative simulated nuclei (blue) containing ecDNA molecules (cyan) generated using different combinations of simulation parameters. (F) Comparison of the percentage of WT and *LBR*-KO COLO320DM cells containing large ecDNA hubs, quantified by DNA FISH. The threshold for large ecDNA hubs was defined as the top 2% of ecDNA FISH objects by area in WT cells. Bars represent the mean of two replicates, and dots indicate individual replicates. (G) Comparison of ecDNA copy number, calculated from Nanopore whole-genome sequencing, between WT and *LBR*-KO COLO320DM cells. Each dot represents an individual replicate (n=3). Error bars indicating standard error of the mean. n.s. represents the two-sided Student’s t-test *P* value > 0.05. (H) Expression levels of *LBR* in TCGA samples stratified by ecDNA status using AmpliconArchitect. The box plots show the expression levels of *LBR* in ecDNA+ and ecDNA-samples. Boxes represent the IQR with center lines indicating the median; whiskers extend to 1.5× IQR. Dots represent individual samples. *P* values were calculated using a two-sided Mann–Whitney U test. TPM, Transcripts Per Million. (I) Expression levels of genes encoding H3K9me3 methyltransferases and HP1 in TCGA samples stratified by ecDNA status. The box plots show the expression levels of genes encoding H3K9me3 methyltransferases and HP1 in ecDNA+ and ecDNA-samples. Boxes represent the IQR with center lines indicating the median; whiskers extend to 1.5× IQR. Dots represent individual samples. *P* values were calculated using a two-sided Mann–Whitney U test. TPM, Transcripts Per Million.

**Figure S8.**
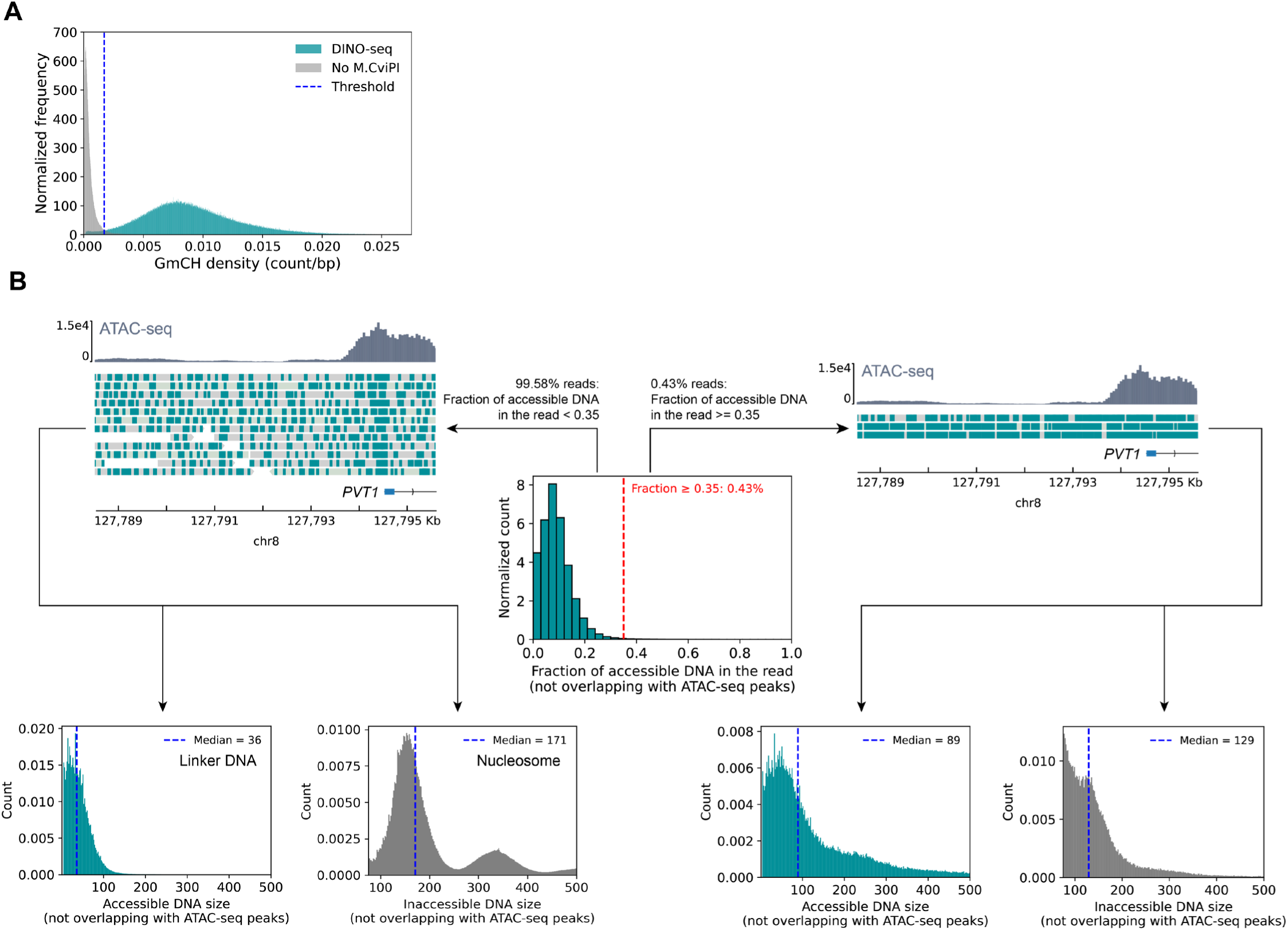
DINO-seq read filtering, related to Methods (A) Comparison of GmCH density (i.e., the number of GmCH modifications on the read divided by aligned read length) of individual reads from DINO-seq with a negative control experiment (i.e., DINO-seq without adding M.CviPI). The blue dashed line indicates the intersection point between the two normalized histograms, representing the threshold that best separates true accessibility-dependent GmCH signal from background. (B) Filtering reads with abnormally high GmCH density. Because the footprint of a nucleosome is ∼146 bp and the linker DNA between adjacent nucleosomes typically spans 0–80 bp^23^, the maximum expected fraction of accessible DNA on a read that do not overlap with ATAC-seq peaks (i.e., composed only of linker DNA) is approximately 80 / (146 + 80) = 0.35. Reads with an apparent accessible fraction above this threshold likely represent aberrant molecules rather than true chromatin features. A small population of such highly accessible reads has also been noted in previously reported single-molecule footprinting assays and is thought to arise from dead or damaged cells^62^. Consistent with this interpretation, reads with abnormally high GmCH levels fail to exhibit the characteristic nucleosome-footprint pattern (shown by the absence of periodic inaccessible-DNA lengths), whereas reads with accessible-DNA fractions below 0.35 display clear nucleosome footprint signatures and typical linker DNA length.

